# “Mitotic” kinesin-5 is a dynamic brake for axonal growth

**DOI:** 10.1101/2024.09.12.612721

**Authors:** Wen Lu, Brad S. Lee, Helen Xue Ying Deng, Margot Lakonishok, Enrique Martin-Blanco, Vladimir I. Gelfand

## Abstract

During neuronal development, neurons undergo significant microtubule reorganization to shape axons and dendrites, establishing the framework for efficient wiring of the nervous system. Previous studies from our laboratory demonstrated the key role of kinesin-1 in driving microtubule-microtubule sliding, which provides the mechanical forces necessary for early axon outgrowth and regeneration in *Drosophila melanogaster*. In this study, we reveal the critical role of kinesin-5, a mitotic motor, in modulating the development of postmitotic neurons.

Kinesin-5, a conserved homotetrameric motor, typically functions in mitosis by sliding antiparallel microtubules apart in the spindle. Here, we demonstrate that the *Drosophila* kinesin-5 homolog, Klp61F, is expressed in larval brain neurons, with high levels in ventral nerve cord (VNC) neurons. Knockdown of *Klp61F* using a pan-neuronal driver leads to severe locomotion defects and complete lethality in adult flies, mainly due to the absence of kinesin-5 in VNC motor neurons during early larval development. *Klp61F* depletion results in significant axon growth defects, both in cultured and *in vivo* neurons. By imaging individual microtubules, we observe a significant increase in microtubule motility, and excessive penetration of microtubules into the axon growth cone in *Klp61F*-depleted neurons. Adult lethality and axon growth defects are fully rescued by a chimeric human-*Drosophila* kinesin-5 motor, which accumulates at the axon tips, suggesting a conserved role of kinesin-5 in neuronal development.

Altogether, our findings show that at the growth cone, kinesin-5 acts as a brake on kinesin-1-driven microtubule sliding, preventing premature microtubule entry into the growth cone. This regulatory role of kinesin-5 is essential for precise axon pathfinding during nervous system development.

## Introduction

Neurons are highly polarized cells that play a central role in the functioning of the entire animal body. They possess specialized compartments —axons and dendrites—critical for the precise transmission and reception of electrical signals. These elongated structures require a well-organized cellular framework, where microtubules are distinctly arranged to support selective transport, ensuring the proper delivery of essential cargoes to both axons and dendrites [1-3].

Molecular motors are protein machines powered by ATP hydrolysis that travel along cytoskeletal filaments in a directed manner. Beyond their role in transporting cargo, several microtubule-based molecular motors and their regulators are essential for building the microtubule network that supports neuron structure and function. Both the plus-end-directed motor kinesin-1 and the minus-end-directed motor cytoplasmic dynein are involved in microtubule organization during neurite outgrowth [4]. Kinesin-1, in particular, drives the sliding of microtubules apart using its ATP-dependent motor domain and an ATP-independent microtubule-binding site located at its C-terminal tail [5]. This kinesin-1-driven microtubule sliding powers cellular shape changes [6] and is essential for initial neurite outgrowth [7], axonal and dendritic extension [5], and axonal regeneration [8].

Interestingly, kinesin-1-driven microtubule sliding relies on the kinesin-1 heavy chain (KHC) but is independent of the kinesin-1 light chain (KLC) [5, 6]. KHC-driven microtubule sliding is further regulated by other molecules. For instance, MAP7/Ensconsin is required for microtubule sliding via activation of the KHC motor [9]. In contrast, the microtubule motor kinesin-6/MKLP1/Pavarotti, known as a component of the centralspindlin complex, crosslinks microtubules to inhibit microtubule sliding [10]. This inhibition is controlled by phosphorylation of kinesin-6/MKLP1/Pavarotti by the Ser/Thr kinase Tricornered (Trc), which modulates the ability of kinesin-6/MKLP1/Pavarotti to bind microtubules [11].

In addition to kinesin-6, another microtubule motor, kinesin-5, has been implicated in microtubule organization during neuronal development [12-19]. Kinesin-5 forms a bipolar homotetramer and plays a crucial role in mitotic spindle formation by sliding antiparallel microtubules apart between spindle poles [20]. Kinesin-5 homologs include *A. nidulans* BimC, *S. pombe* Cut7, *S. cerevisiae* Cin8 and Kip1, *Drosophila* Klp61F, *Xenopus* Eg5, mouse KIF11, and human HsEg5 [21]. The homotetrameric structure of kinesin-5 relies on the central bipolar assembly (BASS) domain [22, 23], and this bipolar conformation is essential for the formation of a bipolar mitotic spindle by sliding antiparallel microtubules and separating spindle poles [24] (Figure 1A). Inhibition of kinesin-5 in mitotic cells leads to a classic monopolar spindle phenotype, where the two spindle poles fuse, and the two half spindles merge [25].

**Figure 1.**
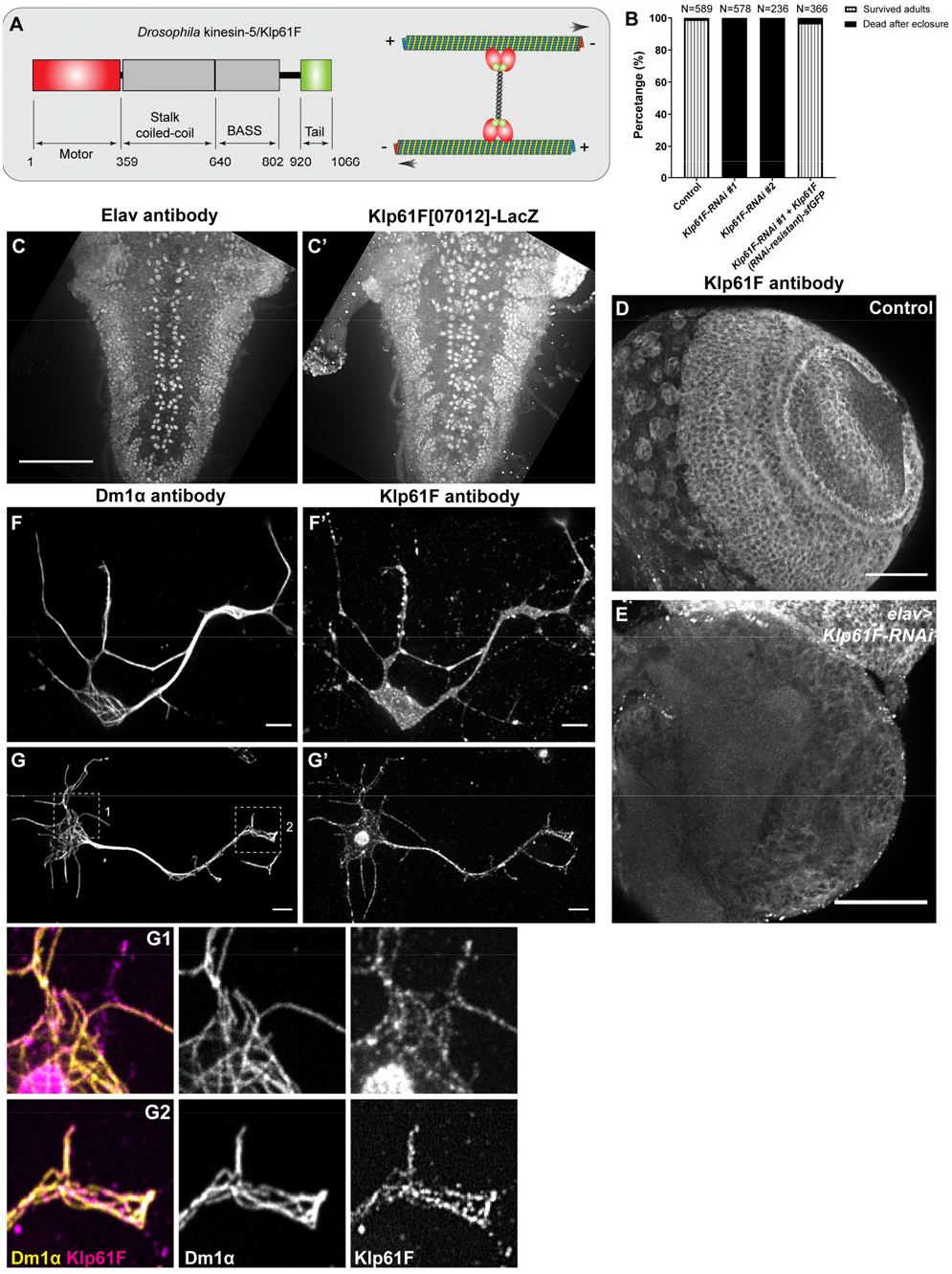
Kinesin-5/Klp61F is essential for neuronal function. **(A)** A schematic illustration of the molecular structure of the *Drosophila* kinesin-5 Klp61F. **(B)** Survival percentage of eclosed adults in control (*elavP>tdEOS-atub84B*), pan-neuronal knockdown of *Klp61F* (*elavP>Klp61F-RNAi #1*, and *elavP>Klp61F-RNAi #2*), and pan-neuronal rescue of *Klp61F* depletion with RNAi-resistant full-length Klp61F (*elavP>Klp61F-RNAi #1+ Klp61F (RNAi-resistant)-sfGFP*). See also Video 1. **(C-C’)** A maximum projection of Z-stacks from the entire 3^rd^ instar larval VNC of a Klp61F enhancer trap line, *Klp61F[07012]-LacZ*, stained with anti-Elav (C) and anti-βGal (C’) antibodies. Scale bar, 50 μm. See also Video 2. **(D-E)** A maximum projection of 2.5 μm slices of a 3^rd^ instar larval control brain (D) and a neuronal *Klp61F*-depleted brain (E) stained with anti-Klp61F antibody, acquired under identical imaging conditions. Scale bars, 50 μm. **(F-G’)** Anti-α-tubulin (DM1α; F and G) and anti-Klp61F (F’ and G’) staining in intact (F-F’) and extracted (G-G’) *Drosophila* larval brain neurons. Scale bars, 5 μm. Zoom-ins of the cell body (G1) and the neurite tip (G2) are shown at the bottom.

In this study, we investigate the role of the *Drosophila* kinesin-5, Klp61F, in postmitotic neurons. Using a combination of classical genetic manipulation and cellular assays, we demonstrate the critical role of Klp61F in neurons. Our findings particularly emphasize its essential function in early motor neurons, where its absence leads to defects in adult locomotion and viability. Additionally, we show that Klp61F regulates axonal growth in both cultured and *in vivo* neurons. Importantly, our findings highlight the significance of Klp61F’s precise localization, especially at axonal tips, where it modulates microtubule penetration into the growth cone. Notably, we reveal that both the microtubule sliding and crosslinking activities of Klp61F are necessary for its function in neurons. In conclusion, we propose that kinesin-5/Klp61F is translocated to the axonal growth cone, where it acts as a “dynamic brake”, regulating axonal outgrowth during neuronal development.

## Results

### Kinesin-5/Klp61F Is Essential for Neuronal Function

Unlike other members of the kinesin superfamily, kinesin5 assembles into a homotetramer through its central BASS domain, located in the stalk region. In this configuration, four motor domains collaborate to slide antiparallel microtubules apart during mitosis (Figure 1A)[22], ensuring the proper formation of the bipolar spindle and accurate chromosome segregation. Kinesin-5 is highly conserved across metazoans, and in *Drosophila*, it is known as Kinesin-like protein at 61F (Klp61F) [26].

We knocked down the fly homologs of four mitotic motors (kinesin-5, kinesin-8, kinesin-11, and kinesin-14) in postmitotic neurons using a pan-neuronal promoter driver, *embryonic lethal abnormal vision (elav)* [27]. The *elav-Gal4* line used in this study was generated by fusing the promoter region of the *Drosophila* gene *elav* with the Gal4 coding region, which was inserted into the 3rd chromosome (hereafter referred to as *elavP-Gal4*) [28]. This line is distinct from the commonly used *elav*^*[C155]*^*-Gal4* line, which is an enhancer trap insertion in the *elav* gene locus [29] known to have expression in non-neuronal tissues [30]. Notably, kinesin-5/Klp61F knockdown by *elavP-Gal4* resulted in severe locomotion defects and 100% lethality in adult flies post-eclosure (Supplemental Figure 1A; Video 1). Two RNAi lines targeting Klp61F produced identical lethal phenotypes, which could be fully rescued by an RNAi-resistant Klp61F full-length transgene (Figure 1B), indicating that the lethality phenotype is caused specifically by *Klp61F* knockdown.

To investigate the cause of the lethality of *elavP>Klp61FRNAi* flies, we first ruled out that the observed effects were due to unintended RNAi knockdown in mitotic tissues. Both mitotic motors kinesin-5/Klp61F and kinesin-13/Klp10A are known to be crucial for fly viability [26, 31]. To confirm the efficiency of our RNAi lines, we performed knockdown of either *Klp61F* or *Klp10A* using a ubiquitous *actin-Gal4*, which resulted in 100% lethality (Supplemental Figure 1A). This outcome confirmed the effectiveness of our RNAi lines in targeting these essential mitotic motors. However, we observed no significant viability defects in *elavP>Klp10A-RNAi* flies (Supplemental Figure 1A), indicating that the lethality seen in *elavP>Klp61F-RNAi* flies was unlikely caused by mitotic defects.

Furthermore, we took advantage of a CDK inhibitor, Roughex (Rux), which disrupts normal cell cycle progression when overexpressed [32-34]. Consistent with previous findings, ubiquitous Rux expression driven by *actin-Gal4* resulted in 100% lethality in adults (Supplemental Figure 1B). However, *elavP-Gal4*-driven Rux expression did not lead to significant lethality (Supplemental Figure 1B), further indicating that the lethality seen in *elavP>Klp61F-RNAi* flies is not due to mitotic defects.

Previous studies have shown that two other *elav-Gal4* lines (*elav*^*[C155]*^*-Gal4* and *elav*.*L2-Gal4*) are transiently expressed in neural progenitor cells, neuroblasts [35]. To rule out the possibility that *elavP>Klp61F-RNAi* affected neuroblast cell division, we examined midline clusters of neurons in the 3rd instar larval ventral nerve cord (VNC) using anti-ELAV staining to label neuronal nuclei. Our data showed that neuronal depletion of *Klp61F* via *elavP-Gal4* did not change the number of neurons in the midline clusters (Supplemental Figure 1C-1D’). These findings suggest that *elavP>Klp61F-RNAi* did not interfere with neuroblast proliferation in the developing nervous system.

Together, these findings indicate that kinesin-5/Klp61F is essential for postmitotic neuron function, and that the lethality observed in *elavP>Klp61F-RNAi* is not caused by mitotic defects.

### Kinesin-5/Klp61F Is Expressed and Localized in Neurons

We examined Klp61F expression using two enhancer trap lines, *Klp61F[07012]-LacZ* and *Klp61F[06345]-LacZ*, in which the P elements carrying the LacZ reporter were inserted 236bp and 1.5kb upstream of the ATG start codon, respectively [26]. Staining for β-galactosidase (βGal) combined with neuronal-specific anti-Elav revealed that Klp61F is highly expressed in larval brain neurons, particularly in the VNC region (Figure 1C-C’; Supplemental Figure 2A-2B’; Videos 2-3).

To further investigate Klp61F localization, we used a rabbit polyclonal antibody specific to Klp61F [36]. Immunostaining showed Klp61F expression in the larval brain (Figure 1D); the staining was significantly reduced in *elav>Klp61F-RNAi* brains (Figure 1E). In cultured neurons dissociated from 3rd instar larval brains, Klp61F immunostaining revealed no strong preference for microtubules (Figure 1F-F’). However, after detergent extraction of soluble proteins, a subpopulation of Klp61F was found on microtubules (Figure 1G-G’), suggesting that a large fraction of Klp61F is inactive and stored in the cytosol.

This finding is further supported by observations in *Drosophila* S2 cells, where a GFP-tagged full-length Klp61F transgene displayed uniform cytosolic localization in intact cells, but complete microtubule decoration after extraction of soluble proteins (Supplemental Figure 2C-C’). Similarly, anti-Klp61F immunostaining in extracted S2 cells showed clear microtubule decoration (Supplemental Figure 2F-F’’).

In summary, Klp61F is expressed and localized in neurons, with its activity is tightly regulated and essential for postmitotic neuronal function.

### Kinesin-5/Klp61F Is Required in Early Motor Neurons for Viability

To investigate the role of Klp61F in neurons, we used a combination of the Gal80 repressor and the UAS-Gal4 system to determine when and where Klp61F is essential for fly viability. Gal4 and Gal80 are yeast-derived proteins with distinct roles: Gal4 is a transcriptional activator that binds to the Upstream Activation Sequence (UAS) to activate gene transcription (such as *Klp61F-RNAi*) [37], whereas Gal80 is a repressor that binds to Gal4’s activation domain, preventing it from interacting with the UAS region [38, 39].

We combined a temperature-sensitive Gal80 under the tubulin promoter (*tubP-Gal80[ts]*) [40] with the pan-neuronal *elavP-Gal4* line. At the permissive temperature (18°C), Gal80[ts] binds Gal4, preventing *Klp61F-RNAi* expression in neurons (RNAi OFF). At the restrictive temperature (29°C), Gal80[ts] is nonfunctional, allowing neuronal *Klp61F-RNAi* expression (RNAi ON). Testing this system, we demonstrated that 18°C fully suppressed the lethality caused by neuronal *Klp61F-RNAi*, while 29°C permitted full penetrance of lethal phenotype in *elav>Klp61F-RNAi* flies (Figure 2A). When freshly eclosed adults raised at 18°C were shifted to 29°C, *Klp61F-RNAi* expression in adult neurons did not increase lethality compared to controls (Figure 2B). Next, we shifted the temperature at the 3^rd^ instar larval stage (29°C→18°C, or 18°C→29°C) to selectively express *Klp61F-RNAi* in neurons before or after the 3^rd^ instar larval stage, respectively. Our results indicated that Klp61F is required in neurons before reaching the 3^rd^ instar larval stage (Figure 2C). Further narrowing the critical window, we found that Klp61F is essential in neurons during early larval development (24-48 hours after egg laying/AEL) (Figure 2C).

**Figure 2.**
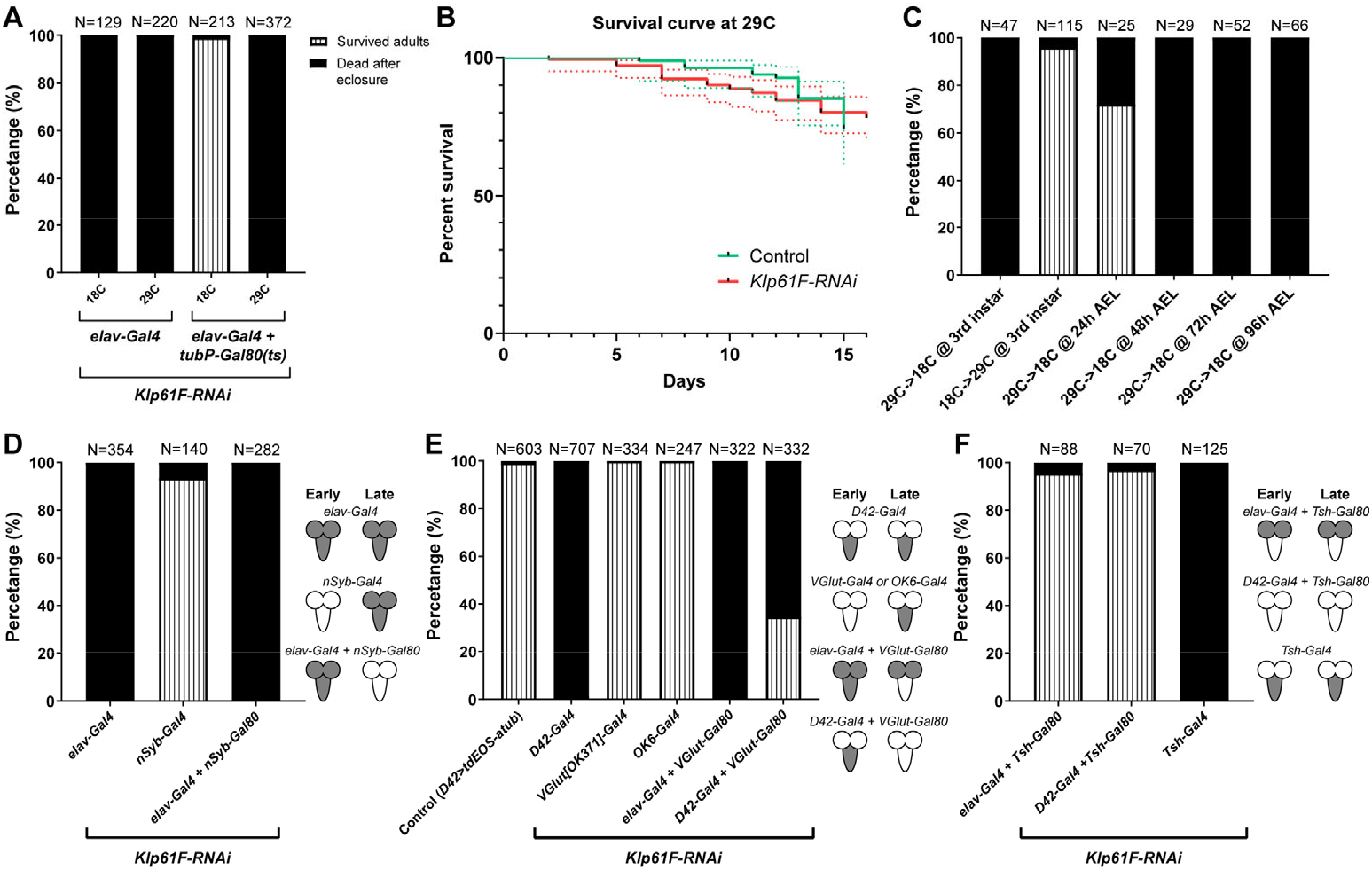
Kinesin-5/Klp61F is required in early motor neurons for viability. **(A)** Efficiency of the Gal4/Gal80(ts) system at permissive (18 °C) and restrictive (29 °C) temperatures. At both 18 °C and 29 °C, *elavP>Klp61F-RNAi* results in 100% lethality in eclosed adults. In contrast, Gal80(ts) suppresses the lethality caused by *elavP>Klp61F-RNAi* at 18 °C but not at 29 °C. **(B)** Adult survival curves of control (*tubP-Gal80(ts)/+; elavP-Gal4/+*) and *Klp61F-RNAi* (*tubP-Gal80(ts)/UAS-Klp61F-RNAi #1; elavP-Gal4/+*). Both samples were raised at the permissive temperature (18 °C, from embryos to adults) and then shifted to the restrictive temperature (29 °C, right after eclosure). N=84 for control and N=141 for *Klp61F-RNAi*. The dotted lines represent the 95% confidence intervals for each survival curve. The P value of the Gehan-Breslow-Wilcoxon test in GraphPrism is 0.8980 (not significant). **(C)** Temperature shifting experiments of *Klp61F-RNAi* flies (*tubP-Gal80(ts)/UAS-Klp61F-RNAi; elavP-Gal4/+*) at different development stages: 29 °C→ 18 °C, at 3^rd^ instar larvae (L3b); 18 °C→29 °C, at 3^rd^ instar larvae (L3b); 29 °C→ 18 °C, at 24 hours, 48 hours, 72 hours and 96 hours after egg laying (AEL). **(D)** Adult survival percentages with both early and late pan-neuronal expression (by *elavP-Gal4*), only late pan-neuronal expression (by *nSyb-Gal4*), and only early pan-neuronal expression (by combining *elavP-Gal4* with *nSyb-Gal80)* of *Klp61F-RNAi*. **(E)** Adult survival percentages with both early and late motor neuron expression (by D42-Gal4), late motor neuron expression (by *VGlut[OK371]-Gal4* or *OK6-Gal4*), or suppression of late motor neuron expression (by combining either *elavP-Gal4* or *D42-Gal4* with *VGlut-Gal80*) of *Klp61F-RNAi*. **(F)** Adult survival percentages with VNC-specific suppression (by combining *elavP-Gal4* or *D42-Gal4* with *Tsh-Gal80*) or VNC-specific expression (*Tsh-Gal4*) of *Klp61F-RNAi*. (D-F) The cartoons on the right illustrate the expression pattern of the *Klp61F-RNAi* (gray) at early and late stages.

Having established the timing of Klp61F’s requirement in neurons, we sought to identify the specific neurons in which Klp61F plays a critical role for fly viability. We first used a later-expressing pan-neuronal Gal4 driver, neuronal *Synaptobrevin (nSyb)-Gal4*, which expresses in mature neurons forming synapse, unlike the early-on *elavP-Gal4*, which is expressed in neurons at all stages, from newly born to fully mature [41]. In contrast to the lethality observed with *elav>Klp61F-RNAi, nSyb>Klp61F-RNAi* caused no significant defects in locomotion or survival (Figure 2D). Furthermore, suppression of *Klp61F-RNAi* expression in mature neurons by *nSyb-Gal80* did not rescue the lethality caused by early neuronal knockdown (*elav>Klp61F-RNAi*) (Figure 2D). This indicates that Klp61F is essential only in early-stage neurons.

Given the severe locomotion defects observed in *Klp61F-RNAi* flies, we hypothesized that Klp61F is particularly critical in motor neurons. Knockdown of Klp61F using the early motor neuron-specific *D42-Gal4* driver [8, 42] phenocopied the lethality seen with *elav>Klp61F-RNAi* (Figure 2E). We excluded the possibility that this lethality was caused by off-target effects in mitotic tissues, as D42-Gal4-driven expression of the CDK inhibitor Rux did not cause significant lethality (Supplemental Figure 1B). Consistent with the idea that Klp61F is required only in early neurons, *Klp61F-RNAi* driven by the more mature motor neuron-specific Gal4 lines, *VGlut*^*[OK371]*^*-Gal4* [43] and *OK6-Gal4* [44], resulted in viable flies (Figure 2E). Moreover, suppression of *Klp61F-RNAi* expression by *VGlut-Gal80* in mature motor neurons did not rescue the lethality driven by *elavP-Gal4* or *D42-Gal4* (Figure 2E).

It has been shown that *D42-Gal4* is ectopically expressed in the peripheral sensory system, particularly in sensory neurons of the body wall [44]. To confirm that the lethality is indeed due to motor neuron dysfunction, we combined *elavP-Gal4* or *D42-Gal4* with *tsh-Gal80*, which inhibits Gal4-driven transcription in the VNC [45]. In both cases, *tsh-Gal80* was able to suppress the lethality caused by *Klp61F-RNAi* (Figure 2F). Additionally, *Klp61F-RNAi* driven by the VNC-specific *tsh-Gal4* [46] resulted in 100% lethality in freshly eclosed adults (Figure 2F). These results indicate that Klp61F is required in VNC motor neurons during early larval stages.

Interestingly, when we assessed 3rd instar larval locomotion by measuring crawling velocity and directionality, we found no significant difference between control and *elav>Klp61F-RNAi* larvae (Supplemental Figure 3A-3B). This suggests that early *Klp61F* knockdown affects motor neurons in the VNC, which continue to develop during metamorphosis and are critical for adult locomotion.

### Kinesin-5/Klp61F Regulates Axonal Growth in Cultured and *in vivo* Neurons

Having mapped the spatial and temporal requirements of Klp61F in neurons, we next investigated the impact of Klp61F depletion on neuronal morphology. We cultured neurons dissociated from 3^rd^ instar larval brains [5, 8, 11, 47, 48] and measured axon length at various time points. In contrast to kinesin-1 knockdown, which significantly reduces axon length [7, 8], Klp61F depletion resulted in longer axons compared to controls (Figure 3A). This suggests that kinesin-5/Klp61F negatively regulates axonal growth.

**Figure 3.**
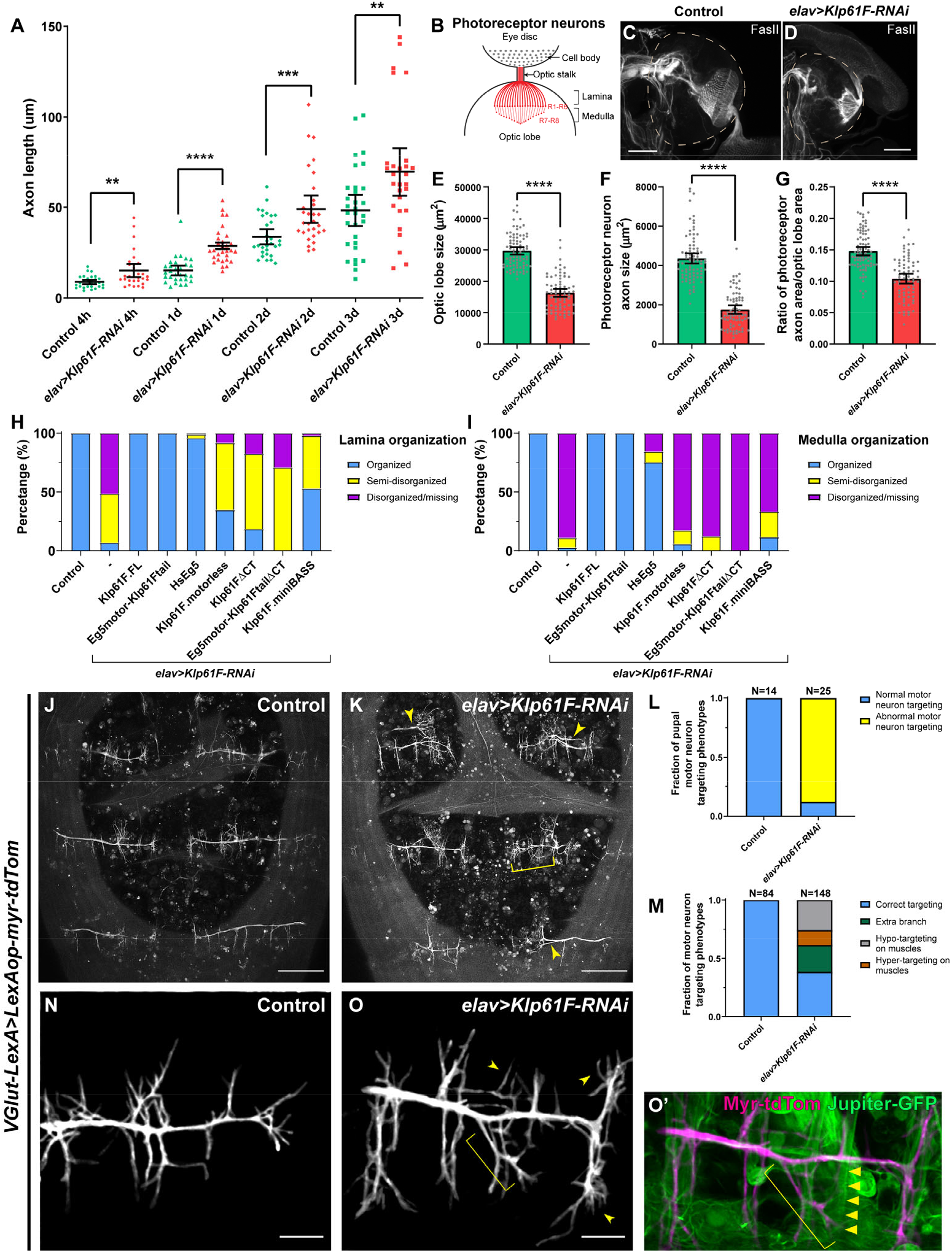
Kinesin-5/Klp61F regulates axonal growth in culture and *in vivo*. **(A)** Axon length measurement in cultured *Drosophila* larval brain neurons of control and neuronal *Klp61F-RNAi*. Neurons were fixed and stained with both anti-α tubulin (DM1α) and anti-Elav antibodies. The length of the longest neurite of each neuron (Elav-positive) was measured as the axon length. Scatter plots with average ± 95 % confidence intervals are shown. Unpaired t-tests with Welch’s correction were performed between control and neuronal *Klp61F-RNAi* samples: 4 hours, p= 0.0018 (**); 1 day, p <0.0001 (****); 2 days, p= 0.0008 (***); 3 days, p=0.0080 (**). **(B)** A schematic illustration of the photoreceptor neurons of *Drosophila* 3^rd^ instar larval eye disc and optic lobe. **(C-D)** Representative examples of photoreceptor neuron axonal targeting in the optic lobes of control (C) and neuronal *Klp61F* depletion (D). Photoreceptor neuron axons were labeled with antiFasciclin II antibody. Dashed lines indicate the position of the optic lobes. Scale bars, 50 μm. **(E-G)** Optic lobe size (E), photoreceptor neuron axon size (F), and the ratio of photoreceptor neuron axon size to the optic lobe size (G) in control and neuronal *Klp61F* depletion. Scatter plots with average ± 95% confidence intervals are shown. Unpaired t-tests with Welch’s correction were performed between control and neuronal *Klp61F-RNAi* samples: optic lobe size, p <0.0001 (****); photoreceptor neuron axon size, p <0.0001 (****); the ratio of photoreceptor neuron axon size to the optic lobe size, p <0.0001 (****). **(H-I)** Summary of lamina (H) and medulla (I) organization phenotypes in control, neuronal *Klp61F-RNAi*, and rescue with RNAi-resistant full-length *Drosophila* Klp61F (Klp61F.FL), chimeric Eg5motor-Klp61Ftail, full-length human Eg5 (HsEg5), and Klp61F.motorless, Klp61FΔCT, chimeric Eg5motor-Klp61FtailΔCT, and Klp61F.miniBASS. Sample sizes for each genotype: control, N=85; *Klp61F-RNAi*, N=72; *Klp61F-RNAi + Klp61F*.*FL*, N=84; *Klp61F-RNAi + Eg5motor-Klp61Ftail*, N=89; *Klp61F-RNAi + HsEg5*, N=97; *Klp61F-RNAi + Klp61F*.*motorless*, N=86; *Klp61F-RNAi* + *Klp61FΔCT*, N=97; *Klp61F-RNAi* + *Eg5motor-Klp61FtailΔCT*, N=24; *Klp61F-RNAi* + *Klp61F*.*miniBASS*, N=51. All samples carried one copy of *elavP-Gal4* and were stained with anti-Fasciclin II antibody to label the photoreceptor neuron axons. **(J-K)** Pupal motor neuron targeting in dorsal abdomen segment A5-A7 in control (J) and in *elav>Klp61F-RNAi* (K) 45-50 hours after pupal formation (APF). Extra branching and hyper-targeting of motor neurons in *elav>Klp61F-RNAi* (K) are shown with arrowheads and a bracket, respectively. Scale bars, 100 μm. **(L-M)** Summary of motor neuron targeting phenotypes, by the numbers of pupae (L) and the numbers of motor neuron groups (M). **(N-O’)** Pupal motor neuron targeting 30 hours APF in control (N) and in *elav>Klp61F-RNAi* (O). Extra (arrowheads) and mistargeting (bracket) neurites were seen in *elav>Klp61F-RNAi* compared to control. (O’) In the overlay image of the motor neuron membrane (magenta) and muscles (green, labeled with Jupiter-GFP), an axonal branch (bracket) showed aberrant targeting, deviating from its expected perpendicular orientation relative to the main axon; instead, it originated from an incorrect position and extended obliquely, eventually innervating a distant muscle (indicated by triangles). Scale bars, 20 μm. (J-K) and (N-O’) Motor neurons were labeled with membrane-targeted tdTomato driven by a motor neuron-specific driver (*VGlut-LexA, LexAop-myr-tdTom*); knockdown of *Klp61F* by RNAi was driven independently by the UAS-Gal4 system (*elavP-Gal4, UASp-Klp61F-RNAi*).

To further explore this finding, we examined axon growth *in vivo*, beginning with larval photoreceptor neurons. These neurons extend axons over hundreds of micrometers from the cell bodies in the eye disc to the optic lobes of 3rd instar larvae. The R1-R6 photoreceptors project their axons in the lamina, while the R7 and R8 photoreceptors project further into the medulla (Figure 3B)[49]. In control samples, photoreceptor neurons labeled with anti-Fasciclin II staining displayed a characteristic “umbrella-like” pattern as they projected into the optic lobes (Figure 3C). However, *Klp61F* depletion in neurons severely disrupted this photoreceptor neuron axon pattern, leading to axonal collapse in the lamina and, more prominently, in the medulla (Figure 3D-3I). Instead of extending properly, these photoreceptor axons either terminated prematurely or projected to incorrect locations, indicating severe defects in the axon pathfinding.

Given that our genetic data indicated Klp61F is essential in VNC motor neurons for adult fly locomotion and viability, we further examined motor neuron targeting during pupal stages. Using a motor neuron-specific membrane marker, we observed that neuronal depletion of *Klp61F* caused a range of motor neuron targeting defects in the muscles of the dorsal abdominal segments (Figure 3J-3M). Closer examination at an earlier stage revealed that *Klp61F-RNAi* mutants exhibit extra and mistargeted neurites compared to controls (Figure 3N-3O’), likely leading to the frequent occurrence of extra branches and abnormal hyper-targeting or hypo-targeting to muscles (Figure 3L-3M). These motor neuron mistargeting defects are consistent with the severe locomotion impairments observed in *Klp61F-RNAi* adults after eclosure (Video 1).

### Kinesin-5/Klp61F Accumulates at the Axonal Tips

To further investigate the role of kinesin-5 in neurons, we generated various transgenes to assess their ability to rescue the neuronal knockdown of endogenous *Klp61F* (Figure 4A). The RNAi-resistant full-length Klp61F transgene completely rescued both the lethality (Figure 1B) and the axonal defects observed *in vivo* (Figure 3H-3I).

**Figure 4.**
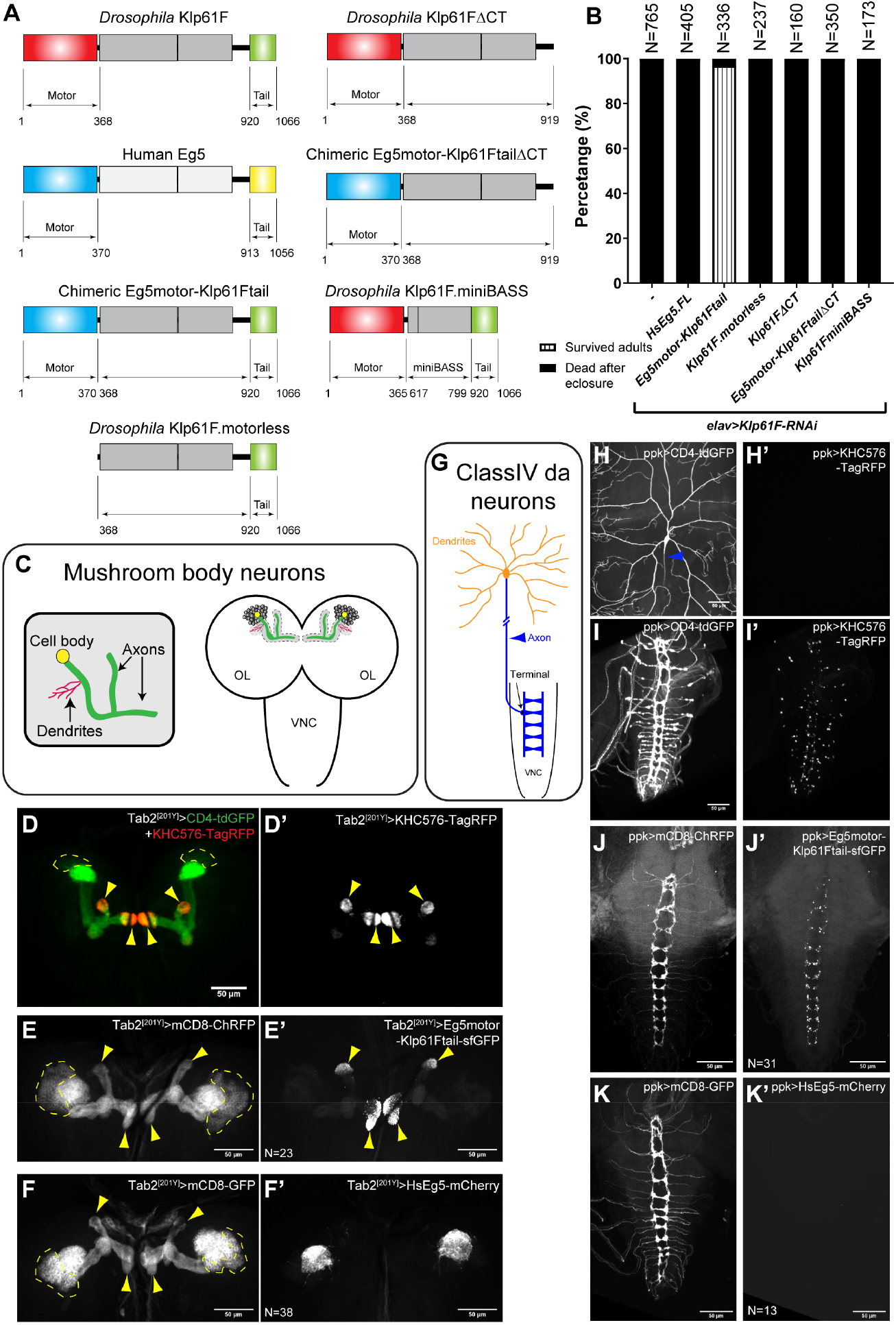
Kinesin-5/Klp61F transgene rescue and neuronal localization. **(A)** Schematic illustrations of transgenes created for *Klp61F-RNAi* rescue experiments. All constructs, except for Human Eg5, carry silent mutations resistant to the *Klp61F-RNAi* (see more details in the “Molecular cloning” section of “Materials and methods”). **(B)** Summary of adult survival percentage in flies expressing neuronal *Klp61F-RNAi* and the listed transgenes. All samples carried one copy of *elavP-Gal4*. See also Video 1. **(C)** Schematic illustration of mushroom body neuron organization. Kenyon cells (the main neurons of the mushroom body) extend the dendrites at the calyx and parallel bundles of axons that are divided into either the α lobe (pointing up) or the β/γ lobes (pointing toward the center). **(D-D’)** Distinct compartments of axons and dendrites in the mushroom body neurons. Axon tips were labeled with a constitutively active kinesin-1 (KHC576). The mushroom body neuron membrane was labeled with CD4-tdGFP, and the dendritic-enrich calyx was labeled with the most concentrated CD4-tdGFP signal. **(E-F’)** Neuronal membrane labeled with mCD8-ChRFP (E-F) and localization of the transgenic chimeric motor (Eg5motor-Klp61Ftail, E’) and full-length human Eg5 (HsEg5, F’) in the mushroom body neurons. (D-F’) All transgene expressions were driven by *Tab2*^*[201Y]*^*-Gal4*. The cell bodies of Kenyon cells are indicated by dashed lines, and their axon tips are indicated by yellow arrowheads. Scale bars, 50 μm. **(G)** Schematic illustration of a class IV da neuron. The da neuron has elaborative dendritic arborization (orange) and extends a single unbranched axon into the VNC (blue). **(H-I)** Membrane labeling (with CD4-tdGFP) of the dendrites (H) and axons (I) of class IV da neurons. The proximal axon from the cell body is indicated by a blue arrowhead, and the axon goes out of focus as it extends deeper beneath the epidermis (H). **(H’-I’)** The constitutively active kinesin-1 (KHC576) motor is concentrated in the axon terminals at the VNC (I’) but is completely absent from the dendrites (H’). **(J-K’)** Neuronal membrane labeled with mCD8-ChRFP (J-K) and localization of the transgenic chimeric motor (Eg5motor-Klp61Ftail, J’) and full-length human Eg5 (HsEg5, K’) in the VNC. The dendritic localization of HsEg5 is shown in Supplemental Figure 5G-G’’. (H-K’) All the transgene expressions were driven by *ppk-Gal4*. Scale bars, 50 μm.

Next, we created a transgene expressing full-length human kinesin-5/Eg5, which successfully rescued mitotic spindle defects in *Drosophila* S2 cells [50](Supplemental Figure 4A-4C). We used the number of gametic ovarioles per ovary as a readout for tissue mitotic index, which depends on proper germline stem cell division and cyst division to populate the germline niche [51]. Inhibition of spindle formation factors, such as XMAP215/Mini spindles (Msps) and kinesin-5/Klp61F in early germline stem cells via *nos-Gal4*^*[VP16]*^ [52, 53] resulted in complete agametic ovaries [54] (Supplemental Figure 4D-4F). Despite its ability to rescue spindle defects in S2 cells, the human Eg5 transgene could not rescue germline stem cell division, resulting in agametic ovaries (Supplemental Figure 4F), nor could it rescue the lethality caused by neuronal *Klp61F* depletion (Figure 4B).

We then generated a chimeric motor consisting of the human Eg5 motor domain and the *Drosophila* Klp61F stalk-tail region (hereafter referred to as Eg5motor-Klp61Ftail) (Figure 4A). This construct not only rescued mitosis *in vivo* (Supplemental Figure 4F), but also rescued adult locomotion and viability following neuronal *Klp61F* knockdown (Figure 4B; Video 1). Furthermore, it fully rescued the photoreceptor neuron axonal targeting defects observed in *Klp61F-RNAi* flies (Figure 3H-3I).

Given that the primary difference between the full-length human Eg5 and the chimeric motor is the *Drosophila* stalk-tail region, we tested whether the motorless stalk-tail construct (hereafter referred to as Klp61F.motorless) was sufficient to rescue the *Klp61F-RNAi* phenotypes. Klp61F.motorless alone failed to rescue the mitotic defects (Supplemental Figure 4F), adult lethality (Figure 4B), or photoreceptor neuron axonal mistargeting (Figure 3H-3I).

Together, these results indicate that both the functional motor domain and the *Drosophila* stalk-tail region are required to fully restore the function of kinesin-5/Klp61F.

sNext, we examined the neuronal localization of these transgenes in two types of highly polarized neurons, mushroom body neurons (Figure 4C) and class IV dendritic arborization (da) neurons (Figure 4G), both of which have well-characterized axonal and dendritic compartments [55, 56]. As axons in both neuron types have uniform plus-ends-out microtubule orientation [57], we used a truncated constitutively active kinesin-1, KHC576-TagRFP [58], to label the axonal tips in mushroom body neurons (Figure 4D-D’) and class IV da neurons (Figure 4H-4I’). The functional chimeric motor Eg5motor-Klp61Ftail was clearly concentrated at the axonal tips in both neuron types (Figure 4E-E’ and 4J-J’), as well as in photoreceptor neurons (Supplemental Figure 5J-5K’).

In contrast, the full-length human Eg5, which could not rescue mitotic and postmitotic defects, was completely absent from axons and predominantly localized to dendrites (Figure 4F-F’ and 4K-K’; Supplemental Figure 5G-5G’’). We also examined the localization of full-length and motorless Klp61F transgenes (Klp61F.FL and Klp61F.motorless, respectively). Full-length Klp61F is predominantly localized to axons with some residual soma localization (Supplemental Figure 5B and 5H), whereas motorless Klp61F was restricted to the cell body and excluded from axons (Supplemental Figure 5C, compared to superfolder GFP (sfGFP) alone in Supplemental Figure 5A; Supplemental Figure 5I).

In summary, we conclude that the functional kinesin-5 transgene is concentrated at the axonal tip, suggesting it plays a critical role there, likely within the axon growth cone.

### Kinesin-5/Klp61F Inhibits Premature Microtubule Penetration into the Cell Periphery

At the axon growth cone, microtubules and actin filaments interact to coordinate axonal guidance, with microtubules extending into the actin-rich peripheral zone to direct axon growth in response to extracellular cues [59, 60]. To better understand the function of kinesin-5 in regulating microtubule interaction with actin cytoskeleton in the growth cone, we employed primary cultures of larval hemocytes [61], as the interaction between microtubules and actin at the periphery of hemocytes closely resembles that of the axonal growth cone (Video 4). Additionally, the large cell size and flat cell shape of hemocytes facilitate high-resolution microscopy. To mimic microtubule penetration into the peripheral region of the growth cone (P-zone) upon exposure to attractive cues, we induced actin fragmentation using Cytochalasin D (CytoD) to the culture. We found that *Klp61F* knockdown resulted in significantly more microtubules penetrating the peripheral area compared to controls, especially after CytoD treatment (Figure 5A-5B; Videos 5-6). We observed similar microtubule over-penetration after immediate kinesin-5 inhibition in *Drosophila* S2 cells, where human Eg5 replaced the endogenous *Klp61F* (Supplemental Figure 6A-6B’; Video 7).

**Figure 5.**
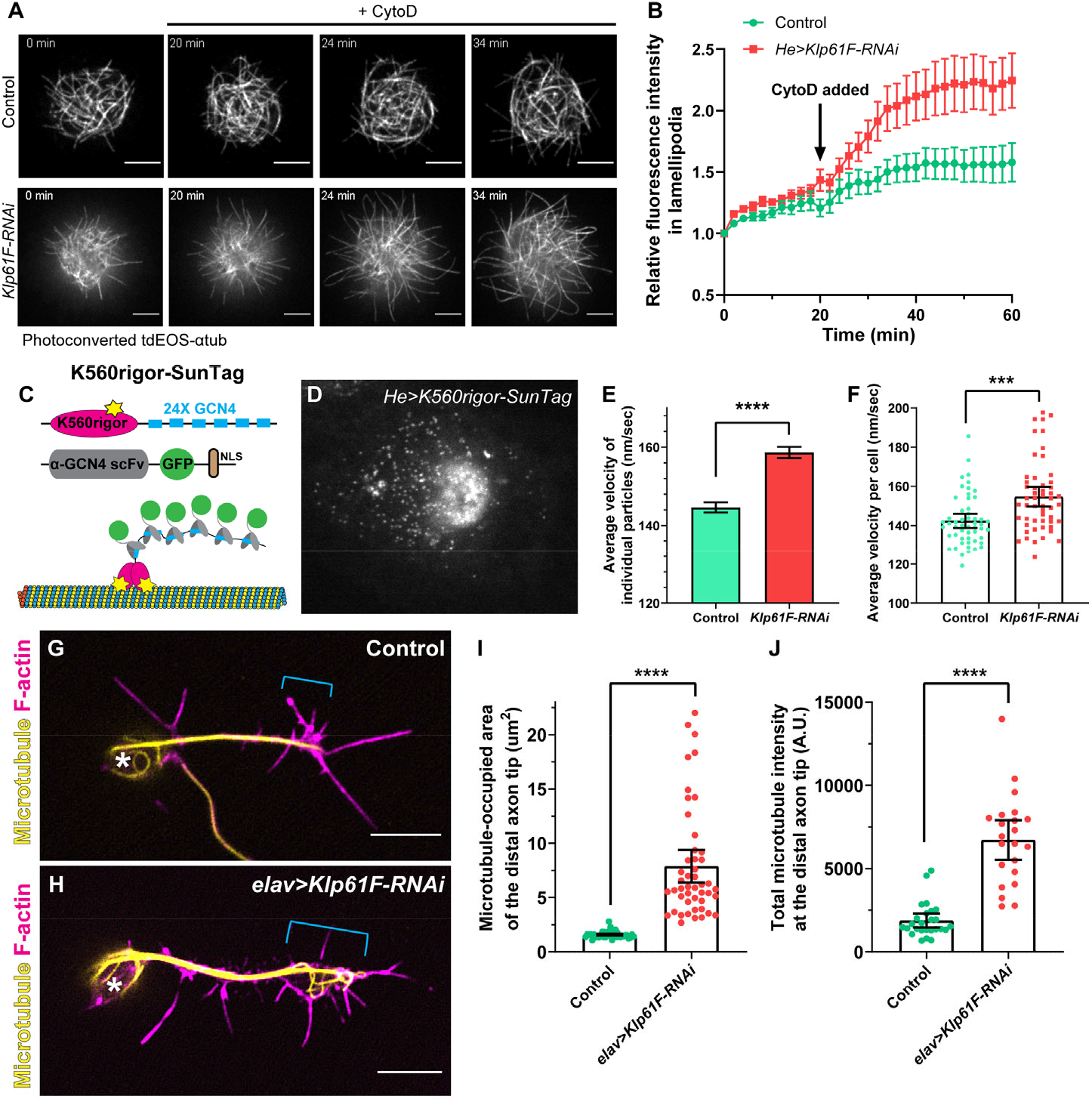
Kinesin-5/Klp61F inhibits microtubule penetration into the periphery of hemocytes and the axonal growth cone of neurons. **(A)** Microtubule penetration before and after actin fragmentation by cytochalasin D (CytoD) in control and *Klp61F*-depleted *Drosophila* primary hemocytes. CytoD was added at 20 minutes after the start of imaging. Microtubules were labeled with α-tubulin 84B tagged with a tandem dimer of EOS (tdEOS-αtub84B), and globally photoconverted from green to red by UV light. Scale bars, 5 μm. See also Videos 5 and 6. **(B)** Quantification of microtubule penetration into the lamellipodia in control and *Klp61F-RNAi* hemocytes. The microtubule intensity in the lamellipodia at 0 minutes was normalized to 1. Data points are shown as average ± SEM. Sample sizes for control and *Klp61F-RNAi* are N=20 and N=24, respectively. **(C)** Schematic illustration of the K560Rigor-SunTag system. Co-expression of (1) a human truncated kinesin-1 motor (K560) carrying a rigor mutation (E236A) with 24 copies of the GCN4 peptide, and (2) a single-chain antibody fragment (scFv) against GCN4 tagged with super-folder GFP (sfGFP) and a nuclear localization signal (NLS), result in bright dots irreversibly attached to microtubules. **(D)** A representative image of a K560Rigor-SunTag-expressing hemocyte. Bright dots outside the nucleus indicate the attachment points of the rigor mutant kinesin-1 motor to microtubules. The movement of these K560Rigor-SunTag dots serves as an indicator of microtubule sliding. **(E-F)** Quantification of K560Rigor-SunTag dot movement in control and *Klp61F-RNAi* hemocytes. Velocities are plotted for all particles tracked (E) or as average velocities per cell (F). The total number of particles tracked is N=8953 from 53 control cells, and N=9857 from 53 *Klp61F-RNAi* cells. The particles were tracked in DiaTrack 3.04. The maximum and minimum velocities were set at 500 nm/sec and 77 nm/sec, respectively. **(G-H)** Distribution of microtubules (by DM1α staining) and F-actin (by phalloidin staining) in control (G) and *elav>Klp61F-RNAi* (H) larval brain neurons 4.5 hours after plating. Asterisks indicate the cell body, and brackets indicate the axon growth cone. Scale bars, 5 μm. Both samples carried one copy of *elavP-Gal4*. **(I-J)** Microtubule distribution in the distal axon tips of control and *elav>Klp61F-RNAi* neurons. The distal axon tip is defined as the distal-most 5 μm of the axon containing microtubules. Microtubule distribution is shown either by the area occupied by microtubules (I) or by the total microtubule fluorescence intensity in the distal tip region (J). Sample sizes for each genotype: (I) control, N=42; *elav>Klp61F-RNAi*, N=47; (J) control, N=26; *elav>Klp61F-RNAi*, N=22. Unpaired t-tests with Welch’s correction were performed between control and *elav>Klp61F-RNAi*: (I), p<0.0001 (****); (J), p<0.0001 (****).

There are two possible explanations for the increased microtubule penetration in *Klp61F-RNAi* cells: increased microtubule polymerization or enhanced microtubule sliding. While previous studies suggested that kinesin-5 may act as a microtubule polymerase, they utilized kinesin-5 in a dimer form rather than its native tetramer configuration [62, 63]. To examine whether *Klp61F* knockdown affects microtubule polymerization, we used the well-characterized microtubule plus-end-binding protein EB1 (EB1-GFP) [54, 64, 65] (Supplemental Figure 6C). Our analysis showed no significant differences in EB1 comet velocity, length, or density between control and *Klp61F-RNAi* cells (Supplemental Figure 6D-6F). Thus, we conclude that increased microtubule polymerization is not the primary reason underlying the enhanced microtubule penetration observed in *Klp61F-RNAi* cells.

We then examined microtubule sliding levels in control and *Klp61F-RNAi* cells directly. To visualize microtubule movement, we employed the K650Rigor-SunTag system, which includes the human kine-sin-1 motor domain with a rigor mutation fused to 24 copies of a GCN4 peptide and a GFP-tagged anti-GCN4 single-chain antibody [66] (Figure 5C). The K560Rigor cannot complete ATP hydrolysis and thus binds irreversibly to microtubules, allowing up to 24 GFP molecules to be recruited to a single microtubule position (Figure 5C). Live imaging of these bright dots provided a direct readout of microtubule movement [67] (Figure 5D). Tracking of these SunTag dots showed that the microtubule sliding rate in *Klp61F-RNAi* cells was significantly higher than in control cells (Figure 5E-5F). These results explain the increased microtubule penetration into the lamellipodia observed in Cy-toD-treated *Klp61F-RNAi* cells (Figure 5A-5B; Videos 5-6).

Given that the functional kinesin-5 transgenes accumulate at the axonal tip, we investigated microtubule organization in the axon growth cones of young cultured larval brain neurons. Consistent with the findings in hemocytes, we observed increased microtubule accumulation in the growth cone following *Klp61F* depletion compared to controls (Figure 5G-J). Based on these results, we propose that kinesin-5 slows microtubule penetration into the growth cone, thereby regulating axon growth.

### Microtubule Sliding and Crosslinking Activities Are Essential for Kinesin-5/Klp61F Function

To determine whether the microtubule sliding function of kinesin-5 is critical in neurons, we created a sliding-deficient version of Klp61F by deleting its C-terminal tail (CT). The kinesin-5 tail binds to the motor domain at the opposite pole in the homotetramer, reducing the motor’s microtubule-stimulated ATP hydrolysis rate, which is necessary for generating sliding forces between microtubules [68]. We deleted the C-terminal tail in both the *Drosophila* full-length and chimeric motors, creating Klp61FΔCT and Eg5motor-Klp61FtailΔCT, respectively (Figure 4A). Although both constructs localized properly to axons, similar to the CT-containing transgenes (Supplemental Figure 5D-5E, compared to Supplemental Figure 5B and Figure 4E’), neither of them was able to restore adult viability (Figure 4B) or photoreceptor axon targeting (Figure 3H-3I) in the neuronal *Klp61F* knockdown flies. This indicates that the microtubule sliding ability, while not required for proper localization at the axon tip, is essential for kinesin-5’s function in neurons.

In addition, we generated a “petite” version of Klp61F by adopting a minimal kinesin-5 tetramer (Klp61F.miniBASS, Figure 4A) [23]. The kinesin-5 miniBASS motors are approximately 38 nm in length, half the length of the native kinesin-5 [23]. This Klp61F.miniBASS construct is less efficient at crosslinking and aligning microtubules due to its shorter, stiffer tetramer BASS filament [23]. Although Klp61F.miniBASS was correctly localized to axon regions in mushroom body neurons (Supplemental Figure 5F), it was unable to rescue adult lethality (Figure 4B) or the photoreceptor axon targeting defects in the optic lobes (Figure 3H-3I) caused by neuronal *Klp61F-RNAi* knockdown. Although this construct is capable of sliding microtubules more efficiently than full-length motors after having crosslinked and aligned microtubules [23], its failure to rescue the phenotypes suggests that microtubule alignment is a prerequisite for the kinesin-5’s sliding function within the neurons.

In conclusion, both the microtubule crosslinking and sliding activities of kinesin-5 are essential for its proper function in neurons.

## Discussion

Kinesin-5 slides antiparallel microtubules within bipolar mitotic spindles, a function essential for cell division. In this study, we demonstrate that beyond its well-known mitotic role, kinesin-5 also plays a crucial role in postmitotic neurons in *Drosophila melanogaste*r: (1) it is expressed and localized in neurons (Figure 1); (2) its function in early motor neurons of the VNC during early larval stages is essential for adult locomotion and viability (Figure 2); (3) it regulates axonal growth both in both cultured and *in vivo* neurons (Figure 3); (4) it specifically localizes to axonal tips (Figure 4); and (5) it inhibits premature microtubule penetration into the growth cone by slowing the microtubule sliding rate (Figure 5).

Based on our data, we proposed a multi-step model for kinesin-5 function in neurons (Figure 6): (1) kinesin-5 is transcribed and translated in the neuronal soma, and its motor activity is required for exiting the soma; (2) the decision to enter the axon, but not the dendrites, depends on the kinesin-5 stalk region, not the C-terminal tail; (3) a fully functional tetramer with proper microtubule alignment and sliding abilities is essential at the axonal growth cone.

**Figure 6.**
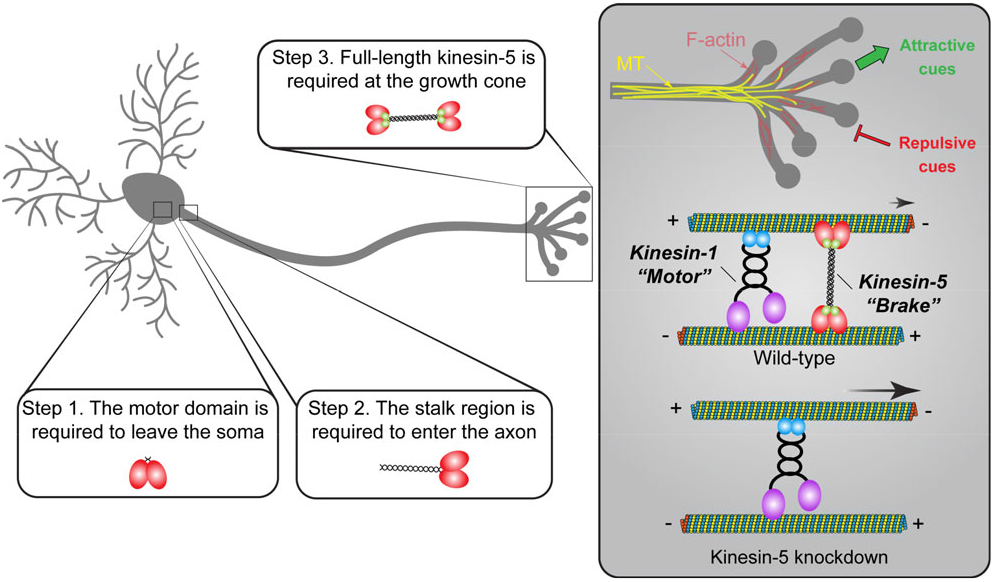
Model of kinesin-5 function in *Drosophila* neurons. Kinesin-5/Klp61F undergoes several critical steps to function properly in neurons: (1) Kinesin-5/Klp61F must leave the soma, a process that requires its motor domain, as motorless Klp61F remains exclusively localized in the neuronal cell body; (2) Kinesin-5/Klp61F must enter the axon, a process dependent on its stalk domain, since the human Eg5 motor lacking the Klp61F stalk domain is mislocalized entirely to the dendrites; (3) The full-length, active tetramer of kinesin-5/Klp61F acts as a dynamic brake on microtubule sliding driven by kinesin-1 at the axon growth cone, playing an essential role in axon pathfinding by responding to both attractive and repulsive cues.

At the growth cone, kinesin-5 slides microtubules at a much slower rate (∼20-40 nm/sec) [69-71] compared to kinesin-1 (∼400-800 nm/sec) [72-74], thereby functioning as a “dynamic brake” for kinesin-1-driven microtubule sliding. We propose that this braking function of kinesin-5 is critical for the correct timing of microtubule penetration into the peripheral zone of the growth cone, allowing the cell to respond properly to both attractive and repulsive cues (Figure 6). In the absence of kinesin-5, faster microtubule sliding results in premature microtubule penetration into the growth cone region, disrupting the tightly controlled process of axon pathfinding (Figure 6).

### The Conservation and Divergence of Kinesin-5 Mitotic Function

While human kinesin-5/Eg5 can rescue mitotic spindle formation in *Drosophila* S2 cells [50], it fails to restore mitotic progression in germline cells depleted of endogenous *Klp61F* (Supplemental Figure 4). This difference between the results from an immortal cell line and *in vivo* data suggests that mitotic regulation is more complex in a living organism. Alignment of human Eg5 and *Drosophila* Klp61F reveals a higher degree of conservation in the motor domains (57% identity, 63% similarity) compared to the stalk-tail region (24% identity, 32% similarity). This suggests that key mitotic regulatory elements may reside in the stalk-tail region, which is consistent with the observation that a chimeric motor (human motor domain and *Drosophila* stalk-tail region) fully restored germline cell division in the absence of endogenous *Klp61F* (Supplemental Figure 4).

One potential explanation is that specific mitotic regulators, such as the kinase Cdk1, phosphorylate the Klp61F tail at T933, which is essential for proper spindle localization and formation [75, 76]. Although human Eg5 has a conserved phosphorylation site at T927, which is similarly targeted by Cdk1 [77, 78], it is possible that *Drosophila* Cdk1 does not properly phosphorylate T927 in human Eg5 (e.g., either hypo-or hyper-phosphorylation), leading to defects in mitotic progression.

On the other hand, replacing the *Drosophila* Klp61F motor domain with the human Eg5 motor domain fully restores the motor function in both mitotic germline cells (Supplemental Figure 4) and postmitotic neurons (Figure 4B; Figure 3H-3I). This highlights the evolutionary conservation between human and *Drosophila* kinesin-5 motor domains. Due to differences in loop5 residues, *Drosophila* Klp61F, unlike *Xenopus* and human Eg5, is insensitive to monastrol and related inhibitors such as S-trityl-L-cysteine (STLC) [79]. By replacing the motor domain with the human version, we successfully “humanized” the flies, making them sensitive to STLC inhibition (Supplemental Figure 4F). This provides an efficient method for pharmacologically inhibiting kinesin-5 activity in flies, facilitating further studies on its role in mitosis and post-mitosis.

### The Regulation of Kinesin-5 Activity by the Stalk-Tail Region

Our data suggest that the *Drosophila* stalk-tail region regulates kinesin-5 microtubule-binding ability, particularly in interphase cells. Kinesin-5/Eg5 is known to localize to spindle microtubules during mitosis but not to interphase microtubules [78]. However, human Eg5 expressed in *Drosophila* S2 interphase cells exhibits strong microtubule decoration. In contrast, both the full-length *Drosophila* Klp61F and the chimeric motor (Eg5motor-Klp61Ftail) are primarily cytoplasmic (Supplemental Figure 2C-2E). This suggests that the *Drosophila* stalk-tail region prevents a significant portion of the motor from binding to interphase microtubules. After detergent extraction, *Drosophila* Klp61F showed microtubule localization in both interphase S2 cells and postmitotic motors (Supplemental Figure 2C’ and 2F-F’’; Figure 1F-1G’), indicating that kinesin-5 retains its microtubule-binding activity in non-mitotic cells, but this activity may be tightly regulated.

In *in vivo* localization assays with highly polarized mushroom body neurons and class IV da neurons, human Eg5 is predominantly localized to the dendrites, whereas the full-length Klp61F and chimeric Eg5motor-Klp61Ftail are concentrated in the axons (Figure 4 and Supplemental Figure 5). Notably, human Eg5 accumulates at the distal dendritic tips in class IV da neurons (Supplemental Figure 5G-G’’), where microtubules display mixed polarity, while the primary dendritic branches contain mostly minus-ends-out microtubules [2]. Kinesin-5 preferentially binds antiparallel microtubules over parallel ones [80], which likely explains the localization of human Eg5 in distal dendrites based on microtubule polarity. In contrast, the functional *Drosophila* full-length Klp61F and chimeric Eg5motor-Klp61Ftail motors show a distinct axonal preference, with minimal localization to the dendrites. This suggests that these functional motors are regulated by unknown mechanisms that prevent them from crosslinking and/or sliding antiparallel microtubules until they reach the target destination in the axons.

Interestingly, while the human Eg5 transgene fails to rescue mitotic progression (Supplemental Figure 4F) and overall viability (Figure 4B), it significantly rescues the photoreceptor neuron targeting defects caused by *Klp61F-RNAi* (Figure 3H-3I). We speculate that this rescue occurs due to the absence of classical dendrites in photoreceptor neurons, which prevents the sequestration of human Eg5 in dendrites. In neurons with well-defined dendrites, such as mushroom body neurons or class IV da neurons, human Eg5 may be sequestered there, limiting its availability in axons. However, without dendritic sequestration in photoreceptor neurons, human Eg5 likely localizes to the axon, where it can carry out its role in regulating axon targeting.

### The Regulation of Kinesin-5 Activity by the Motor Domain

Remarkably, the chimeric motor that rescues both mitotic and postmitotic neuronal phenotypes is more concentrated at the axonal tips of mushroom body neurons and class IV da neurons than the full-length *Drosophila* motor (compared Figure 4E’ and 4J’ to Supplemental Figure 5B and 5H). The chimeric motor co-localizes with a constitutively active kinesin-

1 truncation (KHC576) at the axonal tips, suggesting an intriguing possibility: Klp61F may remain in a processive dimer state rather than a fully assembled tetramer, allowing it to transport itself to the axons. In this scenario, the higher concentration of the chimeric motor at axon tips could be explained by the human motor domain being less regulated than the *Drosophila* motor domain in flies.

In mitotic cells, the *Drosophila* Klp61F motor domain is phosphorylated at three tyrosine residues (Y23, Y152, and Y207) by the tyrosine kinases *Drosophila* Wee1 (dWee1). This phosphorylation is crucial for Klp61F localization and function on microtubules of the mitotic spindle [81]. Of these three residues, Y23 and Y152 are not conserved in human Eg5 (V22 and F154, respectively); notably, the Y152-corresponding site has been replaced by a non-phosphorylatable phenylalanine residue in Eg5. Although dWee1 appears to be an exclusive mitotic regulator, other Tyrosine kinases may phosphorylate these sites in non-mitotic contexts, thereby regulating motor activity. The chimeric motor with the human Eg5 motor domain is likely less tightly regulated than the full-length Klp61F, increasing its likelihood of reaching the axonal tips.

### ATP-Independent Microtubule Binding Site in the Kinesin-5 Tail

In addition to its motor domain, the kinesin-5 tail contains an ATP-independent microtubule-binding site. Motorless constructs have been shown to interact with and bundle microtubules [80, 82, 83]. In kinesin-1, the well-characterized microtubule-binding site in the C-terminal tail consists of a highly conserved stretch of positively charged residues across species that bind to microtubules through electrostatic interactions [4, 5, 67, 84-86]. Unlike kinesin-1, the kinesin-5 tail lacks a conserved cluster of positively charged residues and is instead predicted to be overall negatively charged, which is atypical for interacting with microtubules carrying a strong negative surface charge [83].

Our data from the motorless Klp61F construct show that most of the protein localizes to the cell body and is absent from dendrites or axons (Supplemental Figure 5C). In contrast, soluble GFP is visible throughout the soma, axons, and dendrites (Supplemental Figure 5A). This suggests that the motorless construct is sequestered in the cell body, likely due to the tail’s microtubule binding and bundling activity. This finding indicates that motor activity is essential for kinesin-5 to exit the soma and reach its proper axonal destination.

### Kinesin-5 in Antiparallel Microtubule Sliding in Early Motor Neurons

Here we demonstrate that both the microtubule sliding and crosslinking abilities of kinesin-5 are essential for its function in neurons. Disruption of either ability—whether by deleting the C-terminal tail (which impairs sliding) or by shortening the tetramer filament (which reduces crosslinking)—fails to fully restore kinesin-5’s function in neurons (Figure 4B; Figure 3H-3I). Based on this, we propose a model in which kinesin-5 acts as a “dynamic brake” in the growth cone, aligning microtubules using its native 80 nm-length tetramer and slowly sliding them through coordinated motor activity, modulated by the C-terminal tail. These microtubule crosslinking and slow sliding properties of kinesin-5 decelerate kinesin-1-driven microtubule sliding, thereby preventing premature microtubule penetration into the growth cone during axon pathfinding.

One notable feature is that kinesin-5/Klp61F primarily slides and bundles antiparallel microtubules. Meanwhile, kinesin-1 slides antiparallel microtubules apart and crosslinks parallel microtubules (Figure 6)[4]. However, in *Drosophila* neurons, most axonal microtubules have a uniform plus-ends-out polarity [2], which is less favorable for kinesin-1 sliding and kinesin-5/Klp61F braking activities. Remarkably, previous work from our lab and others has shown that axonal microtubules transition from mixed polarity to uniform polarity during development through a dynein-dependent sorting mechanism [47, 87]. Based on this, we hypothesize that the kinesin-1 (motor) and kinesin-5 (brake) system operates in early axonal growth cones before the establishment of uniform microtubule polarity.

In this study, we found that kinesin-5 is required in VNC motor neurons during early larval stages (Figure 2). However, locomotion in the 3^rd^ instar *Klp61F-RNAi* larvae is indistinguishable from that of control larvae (Supplemental Figure 3), with severe locomotion defects only appearing in early adulthood (Video 1). This suggests that early knockdown of *Klp61F* impacts newly born motor neurons, which are critical for proper locomotion after metamorphosis. This is consistent with our observation that motor neuron targeting to muscles during pupal stages is severely affected by *Klp61F* neuronal depletion (Figure 3J-3R). Altogether, these findings support our hypothesis that kinesin-5 functions as a brake in early axons with mixed-oriented microtubules.

### Novel Aspects of the Kinesin-5 Model

The Baas group has significantly contributed to our understanding of mammalian kinesin-5’s role in neuronal processes, including axonal growth, growth cone turning, dendritic branching, neuronal migration, and axonal regeneration [12-19]. Our findings build upon and complement these discoveries, and while there are many parallels, we identify key distinctions in our conclusions: (1) In our model, kinesin-5 acts as a brake on kinesin-1-driven microtubule sliding, whereas the Baas group suggested that kinesin-5 antagonizes forces generated by cytoplasmic dynein [14]; (2) We observe that kinesin-5 functions specifically in axons, whereas the Baas group found a broader role in regulating dendritic morphology and branching [15]; (3) We did not detect an effect of kinesin-5 on microtubule polymerization, while the Baas group reported increased microtubule polymerization and changes in microtubule orientation following kinesin-5 inhibition [14, 15]; (4) By employing *Drosophila* genetic tools, we were able to examine the role of kinesin-5 *in vivo*, pinpointing its involvement in locomotion and motor neuron targeting. In contrast, the Baas group’s important insights were derived from cell and tissue cultures [12-19].

## Conclusion

Our findings reveal that a “mitotic” motor, kinesin-5, plays an essential role in postmitotic neurons. Kinesin-5 is expressed and localized in neurons, where it acts as a brake on kinesin-1-powered microtubule sliding, thereby preventing premature microtubule penetration into the axon growth cone. Kinesin-5 regulates axon growth both in culture and *in vivo* and is crucial for motor neuron axon targeting, locomotion, and viability. The intricate interplay between kinesin-5 and kinesin-1, functioning as brake and motor in microtubule sliding, provides new insights into axonal development regulation, offering potential avenues for therapeutic intervention in neurodevelopmental disorders.

## Acknowledgements

We thank many colleagues who generously shared reagents: Dr. Thomas J. Maresca (University of Massachusetts), Dr. Gohta Goshima (Nagoya University), Dr. Jawdat Al-Bassam (the University of California, Davis), Dr. Marvin E. Tanenbaum (Hubrecht Institute), Dr. Ronald D. Vale (Janelia Research Campus), and the *Drosophila* Genomics Resource Center (DGRC, supported by NIH grant 2P40OD010949) for DNA constructs; Dr. Chris Q. Doe (University of Oregon), Dr. Christian F. Lehner (University of Zurich), Dr. Elizabeth S. Heckscher and Dr. Edwin L. Ferguson (The University of Chicago), Dr. Melissa M. Rolls (The Pennsylvania State University) and Bloomington *Drosophila* Stock Center (supported by NIH grant P40OD018537) for fly stocks; Dr. Ingrid Brust-Mascher and Dr. Jonathan M. Scholey for anti-Klp61F antibodies. The anti-Elav monoclonal antibody deposited by the Gerald M. Rubin group at Janelia Research Campus, and the anti-Fasciclin II monoclonal antibody deposited by the Corey S. Goodman group at the University of California, Berkeley, were obtained from the Developmental Studies Hybridoma Bank, created by the NICHD of the NIH and maintained at the University of Iowa. We thank all current and past members of the Gelfand laboratory for their support, discussion, and suggestions. This study was supported by the National Institute of General Medical Sciences grant 2R35GM131752 to V.I.G.

## Author contributions

W.L. and V.I.G. designed the research; W.L., B.S.L., H.X.D., M.L., E.M.B., and V.I.G. performed the research; W.L., B.S.L., H.X.D., and E.M.B. analyzed the data; and W.L. and V.I.G. wrote the paper.

## Materials and Methods

### *Drosophila* stocks and husbandry

Fly stocks and crosses were kept on standard cornmeal food (Nutri-Fly Bloomington Formulation, Genesee, Cat # 66–121) supplemented with active dry yeast in a 24 °C incubator. The following flies were used in this study: *elavPGal4* (III, from Dr. Chris Q. Doe)[28]; *UAS-Klp61F-RNAi* #1 (TRiP.GL00441, in the VALIUM22 vector, attP40, Bloomington *Drosophila* Stock center (BDSC) # 35804)[88]; *UAS-Klp61F-RNAi* #2 (TRiP.HMS00552, in the VALIUM20 vector, attP2, BDSC# 33685); *UASp-tdEOS-atub* (II)[7]; *Klp61F[07012]-LacZ* (BDSC# 11710); *Klp61F[06345]-LacZ* (BDSC # 32012); *UAS-Klp67A-RNAi* (TRiP.GL00446, in the VALIUM22 vector, attP40, BDSC #35606); *UAS-Klp10A-RNAi* (TRiP. HMS00920, in the VALIUM20 vector, attP2, BDSC #33963); *UAS-Ncd-RNAi* (TRiP.HMJ22094, in the VALIUM20 vector, attP40, BDSC# 58144); *D42-Gal4* (III, BDSC #8816); *actin5C-Gal4* (III, BDSC# 3954); *UASt-AID-Rux* (III, from Dr. Christian F. Lehner [34]); *tubP-Gal80(ts*) (II, BDSC #7019); *nSyb-Gal4* (III, BDSC# 51635); *nSyb[57C10]-Gal80* (X, from Dr. Elizabeth S. Heckscher); *VGlut[OK371]-Gal4* (II, BDSC# 26160); *OK6-Gal4* (II, BDSC# 64199); *VGlut-Gal80* (II, BDSC# 58448); *Tsh-Gal4, Tsh-Gal80* (II, from Dr. Elizabeth S. Heckscher, [89]); *VGlut-LexA, LexAop-myr-tdTomato* (II)[90]; *Jupiter-GFP* (III, protein trap, ZCL2183, [91]); *Tab2[201Y]-Gal4* (II, BDSC# 4440); *ppk-Gal4* (III, BDSC # 32079); GMR[nina]-Gal4 (II, BDSC# 1104); *UASt-CD4-tdGFP* (8M2, II, BDSC# 35839); *UASt-mCD8-ChRFP* (III, BDSC #27392); *UASt-mCD8-GFP* (II, from Dr. Melissa M.Rolls) [57]; *UASp-sfGFP* (III)[54]; *nos-Gal4[VP16]* (III, from Dr. Edwin L. Ferguson) [52, 53]; *he-gal4* (III, BDSC #8699); *UASp-K560Rigor-SunTag* (*UASp-K560Rigor-24XGCN4, UASp-αGCN4-scFv-sfGFP-NLS*) [66, 67]; *UASt-EB1-GFP* (II) (from Dr. Chris Q. Doe)[8, 47, 57]; *UASt-LifeAct-Rub*y (II, BDSC# 35545).

The following flies (3^rd^ chromosome insertions) were generated in the study by using P-element transformation at BestGene: *UASp-Klp61F*.*FL (RNAi-resistant)-sfGFP, UASp-HsEg5*.*FL-mCherry, UASp-Eg5motor-Klp61Ftail-sfGFP, UASp-Klp61F*.*motorless-sfGFP, UASp-Klp61FΔCT-sfGFP, UASp-Eg5motor-Klp61FtailΔCT-sfGFP* and *UASp-Klp61F*.*miniBASS-sfGFP*.

### Molecular cloning

#### pMT-Klp61F-GFP

The construct was a generous gift from Dr. Gohta Goshima (Nagoya University)[76].

#### pMT-HsEg5-mCherry

The construct was a generous gift from Dr. Thomas J. Maresca (University of Massachusetts, Amherst) [50].

#### pMT-Eg5motor-Klp61Ftail-sfGFP

The construct was generated by inserting human Eg5motor (1-370 residues) and *Drosophila* Klp61F stalk-tail (368-1066 residues) at the SpeI site and sfGFP at the EcoRV site into the pMT vector via Infusion cloning (Takara Bio Inc). Silent mutations were introduced to the region (CDS 2965 nt-2985 nt, 989-995 residues) (CAG-GAGCTGTCCGAAACTGAA→CAAGAACTTAGCGAGACAGAG) by Infusion cloning (Takara Bio Inc) to make the construct resistant to the *UAS-Klp61F-RNAi* #1 line (TRiP.GL00441, attP40, BDSC# 35804).

#### pUASp-Eg5motor-Klp61Ftail-sfGFP

Eg5motor-KLp61Ftail-sfGFP was subcloned from pMT-Eg5motor-Klp61Ftail-sfGFP into pUASp vector via KpnI(5’)/XbaI(3’) sites.

#### pUASp-Klp61F.FL-sfGFP

Klp61F fragment (1-647 residues) was amplified from the pMT-Klp61F-GFP vector and inserted into the pUASp-Eg5motor-Klp61Ftail-sfGFP vector to replace the fragment of human Eg5motor (1-370 residues) + Klp61F stalk (368-647 residues) via KpnI (5’)/SbfI (3’). This construct contains the RNAi-resistant silent mutations (CAAGAACTTAGCGAGACAGAG, corresponding to 989-995 residues).

#### pUASp-HsEg5.FL-mCherry

HsEg5 was subcloned from the pMT-HsEg5-mCherry construct into pUASp by KpnI (5’)/SpeI (3’). mCherry was amplified by PCR from pMT-HsEg5-mCherry and then inserted into the pUASp-HsEg5 vector via the SpeI site.

#### pUASp-Klp61F.motorless-sfGFP

ATG with *Drosophila* Klp61F stalk fragment (368-647 residues) was amplified from pMT-Klp61F-GFP and subcloned into the pUASp-Eg5motor-Klp61Ftail-sfGFP vector to replace the human Eg5 motor via KpnI (5’)/SbfI (3’) to make UASp-Klp61F.motorless (368-1066 residues)-sfGFP. This construct contains the RNAi-resistant silent mutations (CAA-GAACTTAGCGAGACAGAG, corresponding to 989-995 residues). Additionally, a nuclear export signal (NES) (ctgcctcccctggagcgcctgaccctg) was inserted into the pUASp-Klp61.motorless-sfGFP via NotI (5’)/XbaI (3’) to ensure the cytoplasmic localization of the protein.

#### pUASp-Klp61FΔCT-sfGFP

A short piece of oligos (901-919 residues+ linker) (TGAACACCAACGGCAGCAGCTGCAAATTTGCGAGCAAGAGCTTGTACGCTTCAC-TAGTCCAGTGTGGTG) was synthesized and inserted into pUASp-Klp61F-sfGFP vector via SphI(5’)/EcoRI(3’) to create pUASp-Klp61FΔCT (1-919 residues)-sfGFP. This construct contains the RNAi-resistant silent mutations (CAA-GAACTTAGCGAGACAGAG, corresponding to 989-995 residues).

#### pUASp-Eg5motor-Klp61FtailΔCT-sfGFP

Klp61F fragment (901-919 residues)-sfGFP was cut from pUASp-Klp61FΔCT-sfGFP and inserted into pUASp-Eg5mo-tor-Klp61Ftail-sfGFP via SphI(5’)/XbaI(3’). This construct contains the RNAi-re-sistant silent mutations (CAAGAACTTAGCGAGACAGAG, corresponding to 989-995 residues).

#### pUASp-Klp61F.miniBASS-sfGFP

The fragment of Klp61F motor (1-365 residues)-miniBASS (617-799 residues)-partial tail (920-965 residues) was amplified from the Klp61F-minitetramer-withTail-mNeonGreen (a generous gift from Dr. Jawdat Al-Bassam lab) [23] and inserted into the pUASp-Klp61F-sfGFP via KpnI(5’)/BglII(3’) to generate pUASp-Klp61F.miniBASS (1-365 residues + 617-799 residues + 920-1066 residues)-sfGFP. This construct contains the RNAi-resistant silent mutations (CAAGAACTTAGCGAGACAGAG, corresponding to 989-995 residues).

### Larval brain neuron culture

Third instar larval brain neurons were prepared as previously described [5, 8, 11, 48]. Briefly, third-instar larvae were selected and washed with distilled H_2_O, 70% ethanol, and sterile 1× modified dissection saline (9.9 mM 4-(2-hydroxyethyl)-1-piperazineethanesulfonic acid [HEPES], pH 7.5, 137 mM NaCl, 5.4 mM KCl, 0.17 mM NaH_2_PO4, 0.22 mM KH_2_PO4, 3.3 mM glucose, 43.8 mM sucrose). Optic lobes and VNCs were dissected and cleaned in sterile 1× modified dissection saline; tissues were then dissociated in Liberase solution (Roche, Liberase TM Research Grade, 05401119001, diluted in sterile 1× modified dissection saline to final concentration in 0.25 mg/ml) for 1 hour with gentle rotation and pipetting. The neurons were spun down at 300 × g for 5 minutes, and then washed twice in supplemented Schneider’s medium (20% fetal bovine serum, 5 μg/ml insulin, 100 μg/ml penicillin-streptomycin, and 10 μg/ml tetracycline); the neurons were plated onto Concanavalin A (ConA)– coated coverslips and allowed to attach for 20 min before adding the full volume of supplemented Schneider’s medium and incubated for the indicated time before examination.

### Larval brain labeling and immunostaining

Third instar larval brains were fixed and stained as previously described [5, 48]: the larval brains (including optic lobes and VNC) were dissected from 3^rd^ instar larvae in 1X PBS, fixed in 4% EM-grade formaldehyde (Electron Microscopy Sciences 16% Paraformaldehyde Aqueous Solution, Fisher Scientific, Cat# 50-980-487) in 1× PBS +0.1% Triton X-100 for 20 minutes on a rotator at room temperature (24 to 25 °C). The brains were washed with 1× PBTB (1X PBS +0.1% Triton X-100+0.2% BSA) five times for 10 minutes each and blocked in 5% (vol/vol) normal goat serum-containing 1× PBTB for 1 hour at room temperature. The samples were stained with the following primary antibodies at 4 °C overnight: mouse monoclonal anti-Fasciclin II (1D4, DSHB, concentrate, 1:100), mouse monoclonal anti-β-galactosidase antibody (1:250, Promega, Cat# Z3781), rat monoclonal anti-Elav (7E8A10, DSHB, concentrate, 1:100), or rabbit polyclonal anti-Klp61F antibody [36] (1:50). The brains were then washed with 1× PBTB five times for 10 minutes each, and stained with the following secondary antibodies at 10 μg/mL at room temperature for 4 hours: FITC-conjugated anti-mouse secondary antibody (Jackson ImmunoResearch Laboratories, Inc; Cat# 115-095-062; Cat# 715-095-151, minimal cross-reactivity with rat, used with anti-Elav antibody dual staining), TRITC-conjugated anti-rat secondary antibody (Jackson ImmunoResearch Laboratories, Inc; Cat# 712-025-153, minimal cross-reactivity with mouse), or FITC-conjugated anti-rabbit secondary antibody (Jackson ImmunoResearch Laboratories, Inc; Cat# 111-095-003). Finally, the samples were washed with 1× PBTB five times for 10 minutes each before mounting in Mowiol mounting medium.

### Neuron fixation, extraction, and immunostaining

Primary cultured neurons were briefly washed with 1X PBS and 1X Brinkley Renaturing Buffer 80 (BRB80, 80 mM piperazine-N,N’-bis(2-ethanesulfonic acid) [PIPES], 1 mM MgCl2, 1 mM EGTA, pH 6.8) before being fixed with 0.5% glutaraldehyde in 1X BRB80 + 0.1% Triton X-100 for 10 minutes and quenched twice with 2 mg/ml sodium borohydride (NaBH4) in 1X PBS with 10 minutes each. Cells were washed and blocked in Wash Buffer (1X TBS+1% BSA+0.1% TritonX-100) three times for 10 minutes each and then incubated with primary antibodies in Wash Buffer (mouse anti-tubulin DM1α, 1:200; rabbit anti-Klp61F, 1:100; or rat anti-ELAV, 1:100) for 45 minutes. Coverslips were washed in Wash Buffer three times for 10 minutes each and incubated with FITC-conjugated anti-mouse (1:200; Jackson ImmunoResearch Laboratories, Inc; Cat# 115-095-062) and TRITC-conjugated anti-rabbit (1:200; Jackson ImmunoResearch Laboratories, Inc; Cat#111-025-003) or TRITC-conjugated ant-rat (1:200, Jackson ImmunoResearch Laboratories, Inc; Cat# 712-025-153) secondary antibodies in Wash Buffer for 45 minutes. Coverslips were then washed three more times in Wash Buffer 10 minutes each, followed by three quick washes in 1X PBS before mounting in Mowiol mounting medium. For extraction, prior to fixation, cells were incubated in 0.5% Triton X-100 in 1X BRB80 for 1 minute before switching to the fixation buffer. For actin labeling, rhodamine-conjugated or Alexa Fluor 633-conjugated phalloidin (0.2 μg/mL) was included in the secondary antibody staining solution.

### Larval crawling assay

Third instar larvae were washed in dH_2_O for 10 seconds. The larvae were then transferred by a soft brush to the center of a petri dish containing 2% agar supplemented with 25% apple juice. They were allowed to crawl until they reached the edge of the petri dish. ImageJ was used to manually track the larvae and measure both their velocity and directionality.

### Pupal motor neuron imaging

White pupae were handpicked from the vials as 0 hours after pupal formation (APF) and kept in a moisture chamber until the indicated time. Pupal cases were carefully removed on double-sided tape, and 45-50 hours APF pupae were placed on a glass-bottom dish with the dorsal side facing the glass. A small drop of halocarbon oil mix (a 3:1 mix of halocarbon oil 700 and halocarbon oil 27 by volume) was applied between the pupae and the glass. Z-stack images of motor neurons in the dorsal abdomen segments A5-A7 were acquired using an inverted Nikon A1plus scanning confocal microscope with a GaAsP detector and a 20× 0.75 N.A. lens, using Galvano scanning with steps of 0.975 μm. A standard-deviation Z projection of the entire Z-stack was generated in FIJI. For earlier stage motor neuron neurite imaging, 30 hours APF pupae were imaged on a Zeiss LSM 980 Airy Confocal Microscope with a Plan Apo 25X 0.8 N.A. lens in glycerol. Z-stack images of 68 sections with 0.78 μm voxel depth were acquired. VGlut-positive motor nerves were manually segmented from 25 sections. Averaged intensity projections were generated and smoothed employing Gaussian Blur in FIJI.

### Hemocyte preparation and imaging

Hemocyte cultures were prepared based on a previously described protocol [61] with the following modifications: (1) third instar larvae (10 larvae for each coverslip) were selected, washed with distilled H_2_O, sterilized with 70% ethanol, and washed with sterile 1× PBS; (2) each larva was bled using a pair of forceps in a drop of 100 μl of supplemented Schneider’s medium (20% fetal bovine serum, 5 μg/ml insulin, 100 μg/ml penicillin-streptomycin, and 10 μg/ml tetracycline) on a ConA-coated coverslip; (3) the hemocytes were allowed to attach to the coverslips for 20 minutes before adding the full volume of supplemented Schneider’s medium. Hemocytes expressing EB1-GFP or K560Rigor-SunTag under the hemocyte-specific *he-Gal4* driver were imaged using a Nikon W1 spinning disk confocal microscope (Yokogawa CSU with pinhole size of 50 μm) with a Hamamatsu ORCA-Fusion Digital CMOS Camera and a 100× 1.45 N.A. oil lens at a frame rate of one frame every 2 seconds for 1 min, controlled by Nikon Elements software. EB1-GFP comets and K560Rigor-SunTag dots were tracked in DiaTrack 3.04 as previously described [54, 67].

### Da neuron dendrite imaging

Third-instar larvae were selected and washed with distilled H_2_O, then mounted in 100% glycerol with a dorsal-side-up orientation in a glass sandwich (between a coverslip and a slide with two layers of tape as spacers). For each larva, both left and right dorsal class IV da neurons of segments 4-6 (A1-A3) were imaged on a Nikon W1 spinning disk confocal microscope (Yokogawa CSU with pinhole size of 50 μm) with a Hamamatsu ORCA-Fusion Digital CMOS Camera and a 20× 0.75 N.A. lens, using 0.9 μm/step for z-stacks, controlled by Nikon Elements software.

### S2 cells transfection and RNAi treatment

*Drosophila* S2 cells were maintained in Insect-Xpress medium (Lonza) in a 25°C incubator. Transfections with pMT constructs were performed using Effectene (Qiagen, Cat # 301425) according to the manufacturer’s instructions and induced with 200 μM CuSO4. dsRNA against Klp61F 3’UTR (1-364 nt) was amplified from a Klp61F cDNA template (LD15641; DGRC Stock Number 3647) following a standard *in vitro* transcription protocol. A total of 40 μg dsRNA was added to the cells—20 μg on day 1 and 20 μg on day 3 [5]. S2 cells were then plated on ConA-coated coverslips and immunostained on day 5 [92].

### S2 cell fixation, extraction, and immunostaining

*Drosophila* S2 cells were either fixed with 100% methanol at – 20°C for 10 minutes (for Dm1α and DAPI staining), or with 0.5% glutaraldehyde in 1X PBS for 10 minutes and then quenched twice in 2 mg/ml sodium borohydride (NaBH4) in 1X PBS (for anti-Klp61F + Dm1α double staining) for 10 minutes each at room temperature. Cells were washed and blocked in Wash Buffer (1X TBS+1% BSA+0.1% TritonX-100) three times for 10 minutes each and then incubated with primary antibody in Wash Buffer (mouse anti-tubulin (DM1α), 1:200, and/or rabbit anti-Klp61F, 1:100) at room temperature for 45 minutes. Coverslips were washed in Wash Buffer three times for 10 minutes each and then incubated with FITC-conjugated anti-mouse (1:200; Jackson ImmunoResearch Laboratories, Inc; Cat# 115-095-062) and/or TRITC-conjugated anti-rabbit (1:200; Jackson ImmunoResearch Laboratories, Inc; 111-025-003) secondary antibodies with DAPI (1 μg/mL) in Wash buffer at room temperature for 45 minutes. Coverslips were then washed three times in Wash Buffer for 10 minutes each, followed by three quick washes in 1X PBS before mounting in Mowiol mounting medium. For extraction, prior to fixation, cells were incubated in 0.5% TritonX-100 in 1X BRB80 for 1 minute before switching to the fixation buffer.

## Supplemental Information

**Supplemental Figure 1.**
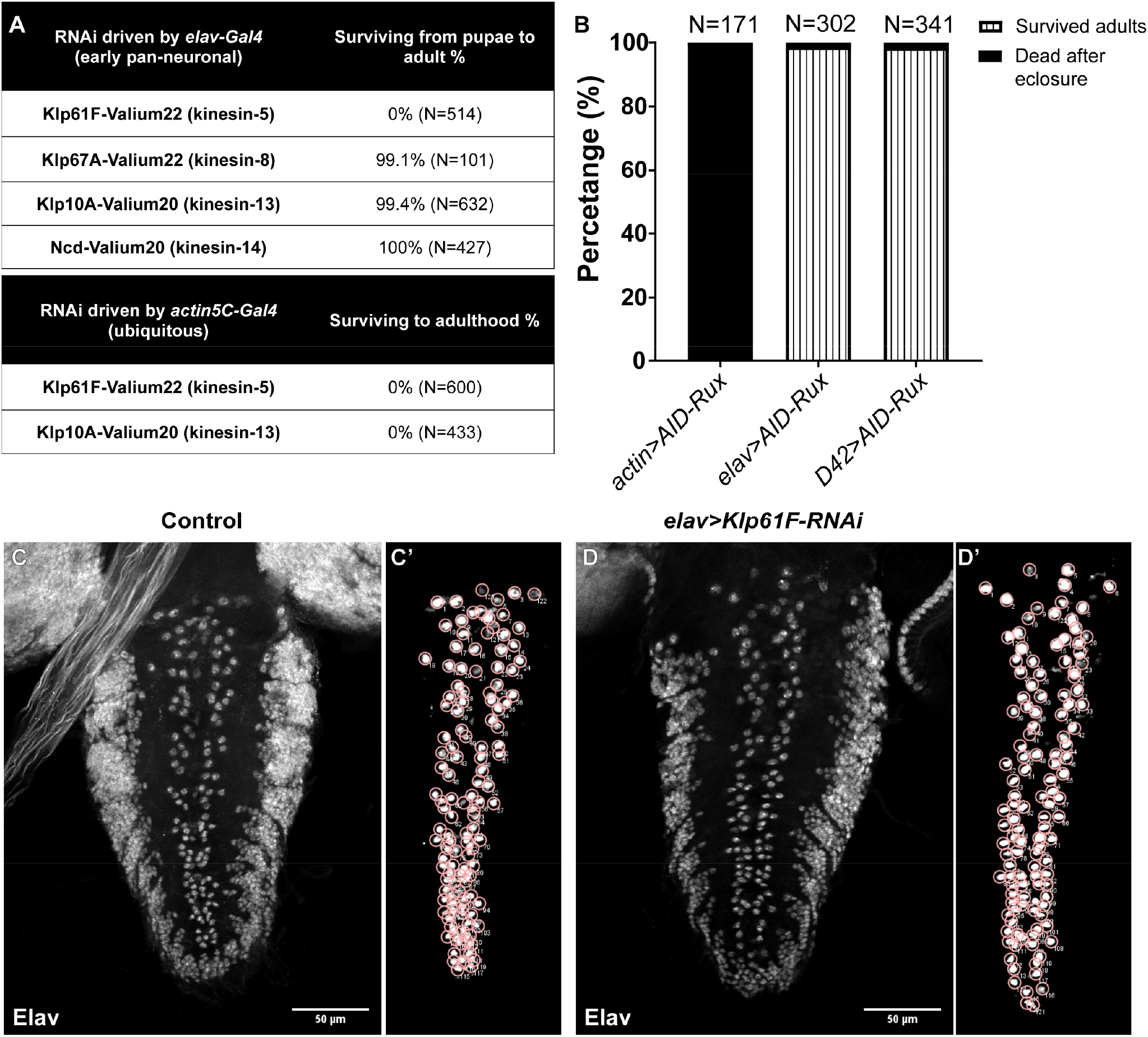
Kinesin-5/Klp61F is required in neurons. **(A)**Summary of adult survival rates following neuronal depletion of four mitotic motors, kinesin-5/Klp61F, kinesin-8/Klp67A, kinesin-13/Klp10A or kinesin-14/Ncd (top), and after ubiquitous depletion of kinesin-5/Klp61F or kinesin-13/Klp10A (bottom). **(B)**Adult survival rates of flies expressing the CDK inhibitor, Roughex (Rux), driven by the ubiquitous *actin5C-Gal4*, pan-neuronal *elavP-Gal4* or pan-motor neuron *D42-Gal4* drivers. **(C-D’)** Neuronal nucleus labeled with anti-ELAV antibody in the 3rd instar larval VNC of control (C) and *elav>Klp61F-RNAi* (D). The manual tracking of the nuclei in the midline clusters of neurons is shown in (C’-D’). Scale bars, 50 μm.

**Supplemental Figure 2.**
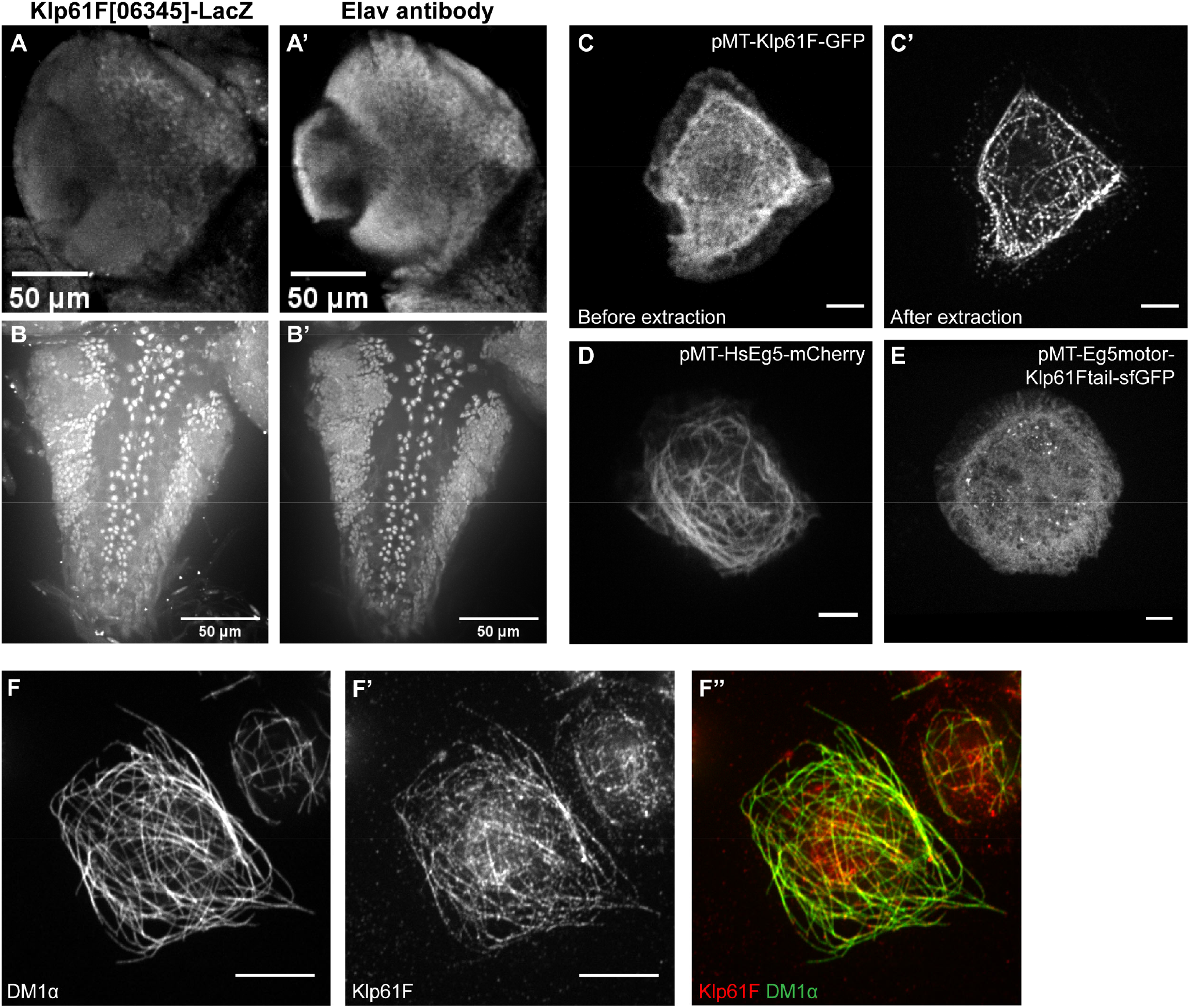
Kinesin-5/Klp61F localization in larval brains and S2 cells. (A-B’) Anti-β-galactosidase staining (A-B) and anti-Elav staining (A’-B’) of the optic lobe (A-A’) and VNC (B-B’) in the Klp61F[06345]-LacZ enhancer trap line. Scale bars, 50μm. See also Video 3. **(C-C’)** Representative images of an S2 cell transfected with pMT-Klp61F-GFP before (C) and after extraction (C’). Scale bars, 5μm. **(D-E)** Representative images of S2 cells transfected with pMT-HsEg5-mCherry (D) and pMT-Eg5motor-Klp61Ftail-sfGFP (E). Scale bars, 5 μm. **(F-F’’)** Anti-α-tubulin (DM1α) staining (F), anti-Klp61F staining (F’), and the overlay of both staining channels (F’’) in extracted S2 cells.

**Supplemental Figure 3.**
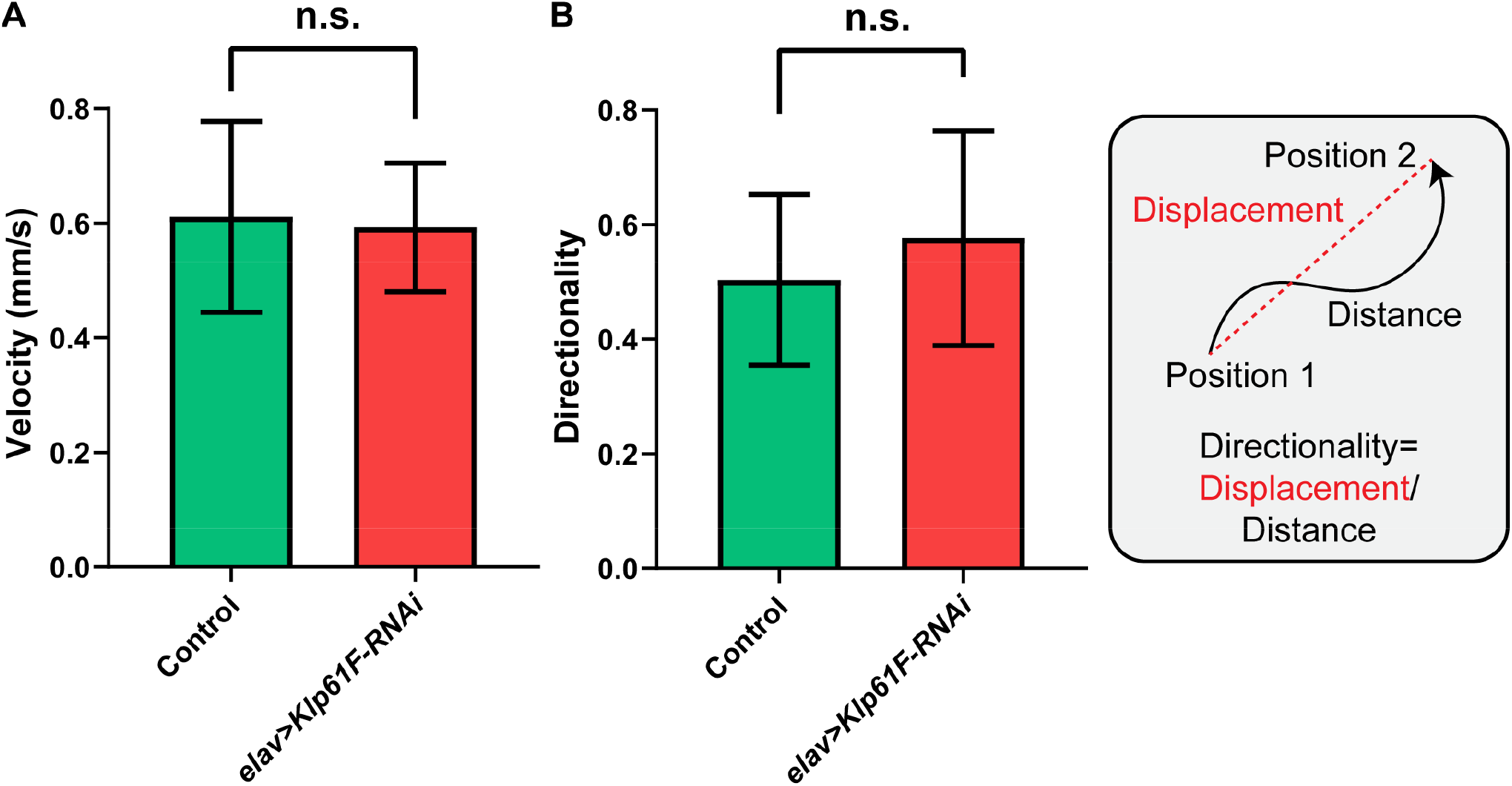
Larval crawling is not affected by neuronal *Klp61F-RNAi*. (A) Velocity and (B) directionality of crawling in 3rd instar larvae with control and neuronal *Klp61F* depletion. Directionality is calculated as the ratio of the displacement to the distance from the center (starting point) to the edge (ending point) of the petri dish. Data are presented as average ± 95% confidence intervals. Sample sizes for control and *Klp61F-RNAi* are N=18 and N=15, respectively. Unpaired t-tests with Welch’s correction were performed between control and *Klp61F-RNAi* samples: Velocity, p= 0.8450 (not significant); Directionality, p=0.5226 (not significant). See “Larval crawling assay” in “Materials and Methods” for more details.

**Supplemental Figure 4.**
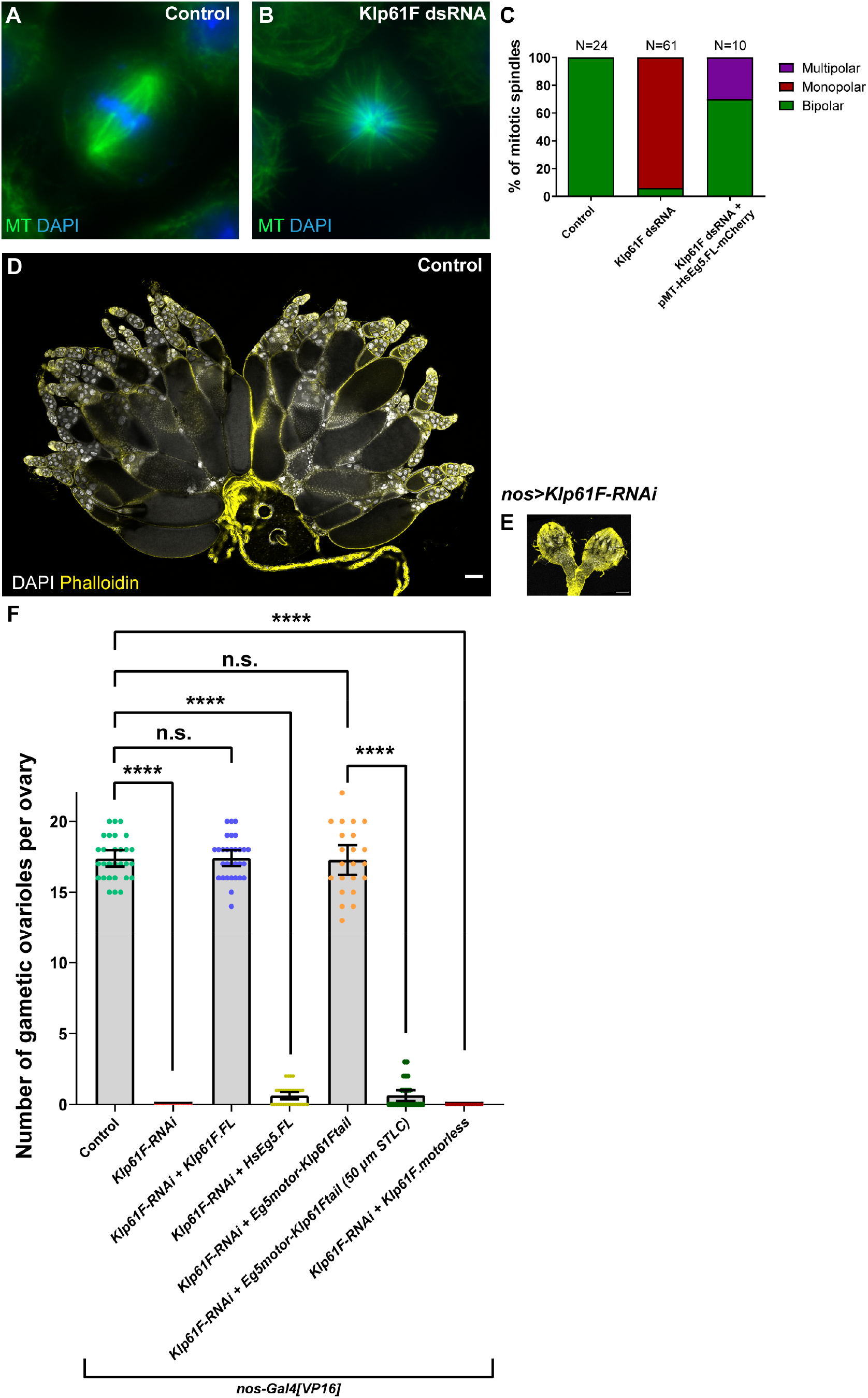
Kinesin-5 transgene rescue in S2 cells and ovaries. **(A-B)** Representative images of a bipolar mitotic spindle in a control S2 cell (A) and a monopolar mitotic spindle in an S2 cell treated with Klp61F dsRNA (B). (C) Percentage of mitotic spindle phenotypes in control, Klp61F dsRNA-treated, and Klp61F dsRNA-treated S2 cells transfected with pMT-HsEg5-mCherry. **(D-E)** Representative images of a pair of control ovaries (D) and a pair of ovaries with *Klp61F* depletion driven by the early germline driver *nos-Gal4*^*[VP16]*^ (E), shown at the same scale. Scale bars, 100 μm. **(F)** Quantification of the number of gametic ovarioles per ovary in listed genotypes. Sample sizes: control, N=29; *Klp61F-RNAi*, N=24; *Klp61F-RNAi + Klp61F*.*FL* (Klp61F.FL rescue), N=30; *Klp61F-RNAi + HsEg5*.*FL* (HsEg5 rescue), N=32; *Klp61F-RNAi + Eg5motor-Klp61Ftail* (chimeric Eg5motor-Klp61Ftail rescue), N=22; *Klp61F-RNAi + Eg5motor-Klp61Ftail* raised with 50 μM STLC (chimeric Eg5motor-Klp61Ftail rescue with STLC), N=27; *Klp61F-RNAi + Klp61F*.*motorless* (Klp61F.motorless rescue), N=28. Unpaired t-tests with Welch’s correction were performed for the following groups: control vs. *Klp61F-RNAi*, p<0.0001 (****); control vs. Klp61F.FL rescue, p=0.9576 (not significant); control vs. HsEg5 rescue, p <0.0001 (****); control vs. chimeric Eg5motor-Klp61Ftail rescue, p= 0.8538 (not significant); control vs. Klp61F.motorless rescue, p <0.0001 (****); chimeric Eg5motor-Klp61Ftail rescue with or without STLC, p <0.0001 (****). Notes: (1) 50 μM STLC was mixed into the cornmeal food where the flies were raised from embryos to adults (no obvious lethality was observed at this concentration of STLC); (2) In the HsEg5 rescue samples, only one or two egg chambers were observed per rescued gametic ovary, often with abnormal numbers of germline cells, indicating poor rescue of mitotic progression.

**Supplemental Figure 5.**
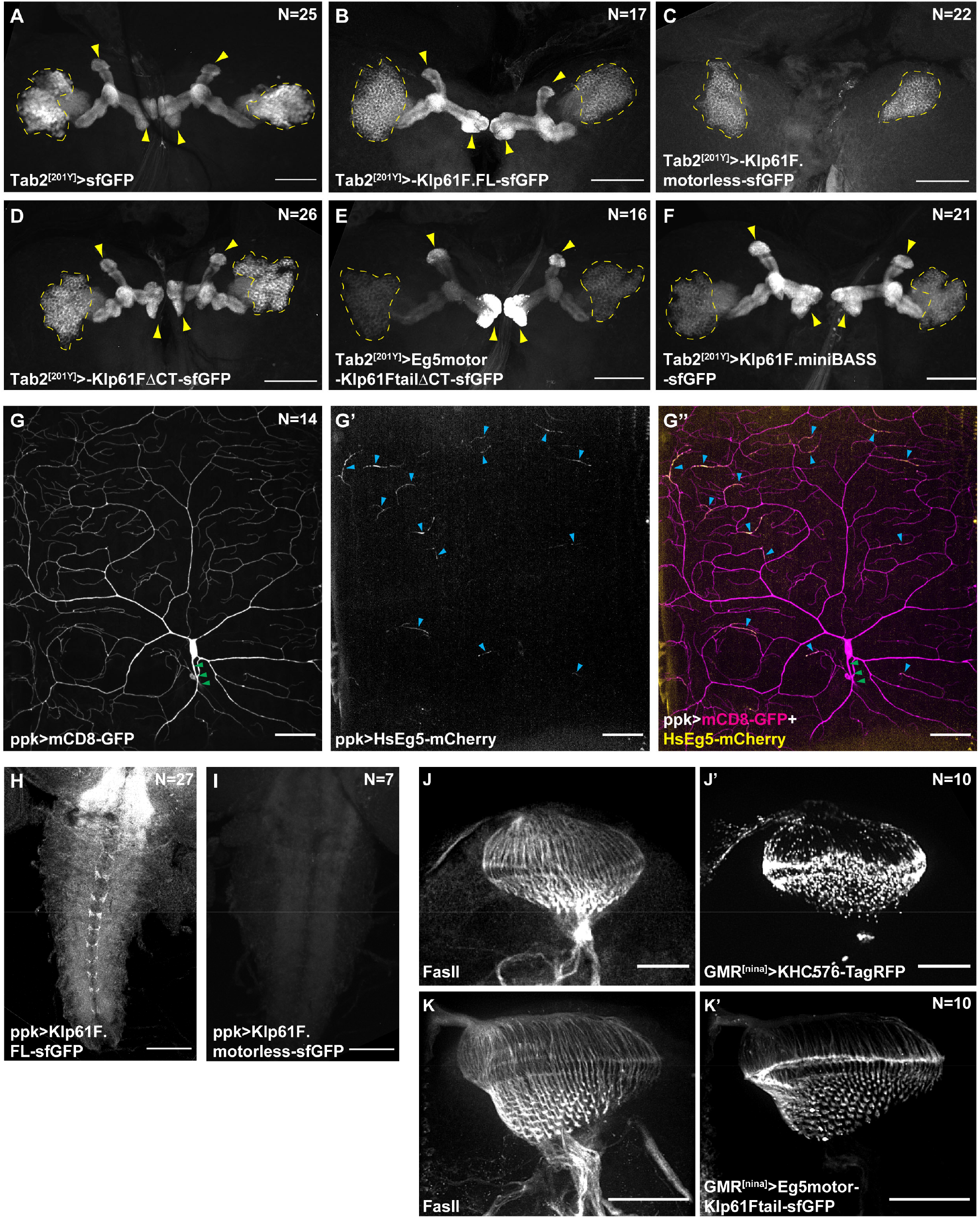
Kinesin-5 transgene localization. **(A-F)** Transgenic expression of soluble sfGFP (A), Klp61F.FL (B), Klp61F.motorless (C), Klp61FΔCT (D), Eg5motor-Klp61FtailΔCT (E), and Klp61F.miniBASS (F) in mushroom body neurons. Details of these transgene constructs are provided in Figure 4A and the “Molecular cloning” section of “Materials and Methods”. All transgene expressions were driven by *Tab2*^*[201Y]*^*-Gal4*. The cell bodies of Kenyon cells are indicated by dashed lines, and their axon tips are indicated by yellow arrowheads. Scale bars, 50 μm. **(G-G’’)** Membrane labeling (mCD8-GFP) and localization of human Eg5 (HsEg5-mCherry) in a class IV da neuron. The proximal axon and the dendritic tips are marked by green and blue arrowheads, respectively. HsEg5 transgenic expression was driven by *ppk-Gal4*. Scale bars, 50 μm. **(H-I)** Transgenic expression of Klp61F.FL (H) and Klp61F.motorless (I) in the axonal terminals of class IV da neurons in the VNC. Both transgene expressions were driven by *ppk-Gal4*. Scale bars, 50 μm. **(J-K’)** Anti-Fasciclin II (FasII) staining (J-K) and localization of either a constitutively active kinesin-1 truncation (KHC576) (J’) or the chimeric Eg5mo-tor-Klp61Ftail transgene (K’). Transgene expression was driven by *GMR*^*[nina]*^*-Gal4*. Scale bars, 50 μm.

**Supplemental Figure 6.**
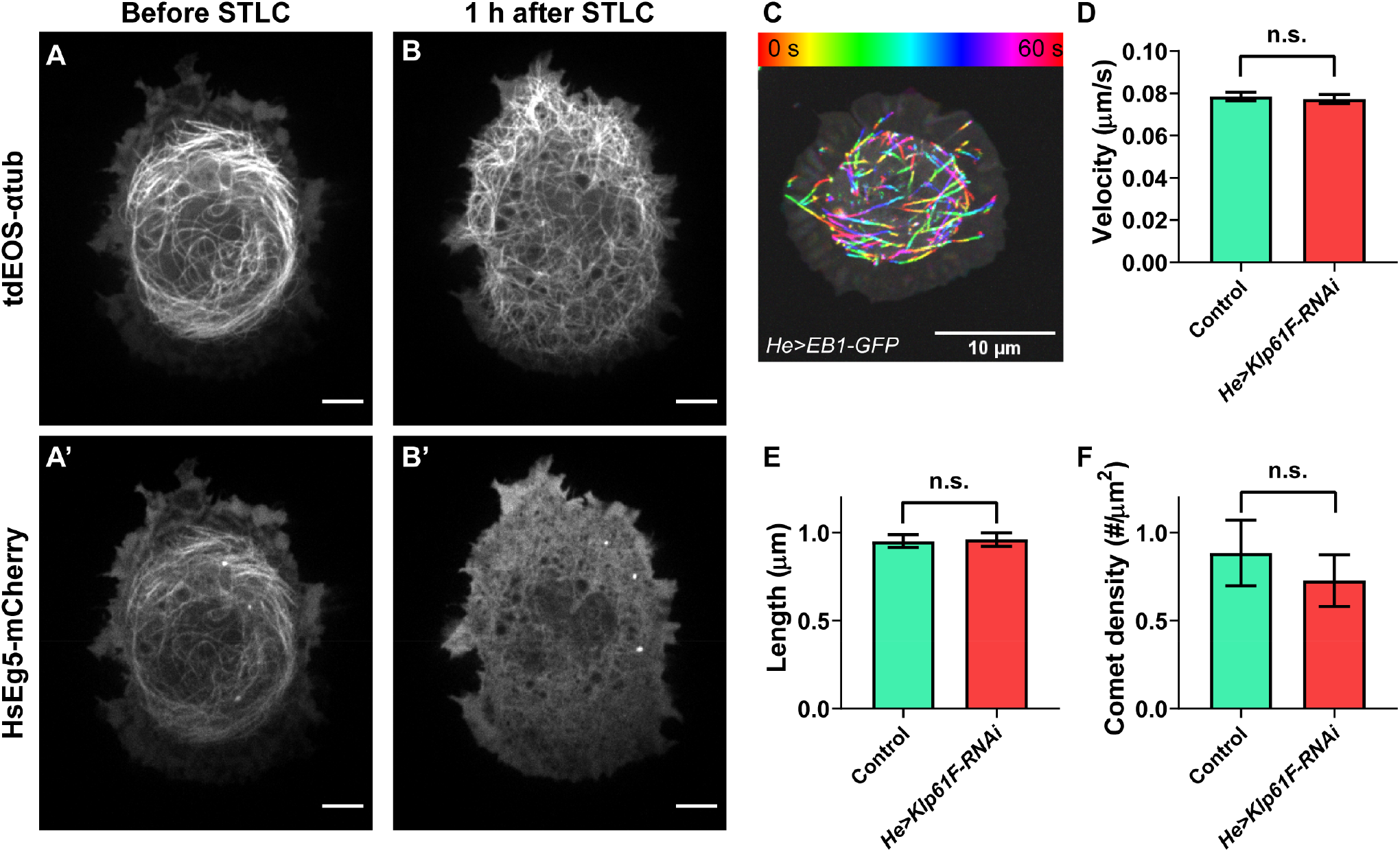
Kinesin-5 prevents microtubule penetration independent of microtubule polymerization. **(A-B’)**Microtubule and human Eg5 localization before and after 2 μM STLC treatment in an S2 cell treated with Klp61F dsRNA. Scale bars, 5 μm. **(C)** A temporal-color hyperstack of EB1-GFP movement in a *Drosophila* primary hemocyte. EB1-GFP expression was driven by *he-Gal4*. Scale bar, 10 μm. **(D-F)** Tracking of EB1 comet velocity (D), length (E), and density (F) in control and *Klp61F-RNAi* hemocytes. Numbers of EB1 comet tracks: control, N=2101; *Klp61F-RNAi*, N=1836. Unpaired t-tests with Welch’s correction were performed between control and *Klp61F-RNAi*: (D) velocity, p=0.4126 (not significant); (E) length, p=0.7163 (not significant); (F) comet density, p=0.1621 (not significant).

## Supplementary Videos

**Video 1. Adult fly climbing assay in control, *Klp61F-RNAi* and the chimeric motor rescue flies**. From left to right: adult flies of control, *elav>Klp61F-RNAi* and *elav>Klp61F-RNAi+Eg5motor-Klp61Ftail-sfGFP*. The video is shown in real-time.

**Video 2. Klp61F[07012]-LacZ and Elav staining in a larval VNC**. Dual staining with anti-Elav (left) and anti-β-Gal (right) in the *Klp61F[07012]-LacZ* enhancer trap line. Scale bar, 50 μm.

**Video 3. Klp61F[06345]-LacZ and Elav staining in a larval brain**. Dual staining with anti-Elav (left) and anti-β-Gal (right) in the *Klp61F[06345]-LacZ* enhancer trap line. Scale bar, 50 μm.

**Video 4. Microtubule movement and actin ruffling in control hemocyte**. Microtubules (labeled with tdEOS-αtub84B, green, without photoconversion) and F-actin (labeled with LifeAct-Ruby, red) driven by *He-Gal4* in a control hemocyte. Scale bar, 5 μm.

**Video 5. Microtubule penetration into the cell periphery in control**. Microtubules (labeled with tdEOS-αtub84B, photoconverted to red) in a control hemocyte before and after 2.5 μM CytoD treatment. Scale bar, 5 μm.

**Video 6. Microtubule penetration into the cell periphery in *Klp61F-RNAi***. Microtubules (labeled with tdEOS-αtub84B, photoconverted to red) in a *He>Klp61F-RNAi* hemocyte before and after 2.5 μM CytoD treatment. Scale bar, 5 μm.

**Video 7. Microtubule penetration into the cell periphery in a Drosophila S2 cell upon STLC treatment**. Microtubules (labeled with tdEOS-αtub84B, green channel, without photoconversion) in a *Drosophila* S2 cell treated with Klp61F dsRNA and ectopically expressing HsEg5-mCherry before and after 2 μM STLC treatment. Scale bar, 5 μm.

